# Lexical frequency and sentence context influence the brain’s response to single words

**DOI:** 10.1101/2020.12.08.416016

**Authors:** Eleanor Huizeling, Sophie Arana, Peter Hagoort, Jan Mathijs Schoffelen

## Abstract

Typical adults read remarkably quickly. Such fast reading is facilitated by brain processes that are sensitive to both word frequency and contextual constraints. It is debated as to whether these attributes have additive or interactive effects on language processing in the brain. We investigated this issue by analysing existing magnetoencephalography data from 99 participants reading *intact and scrambled sentences.* Using a cross-validated model comparison scheme, we found that lexical frequency predicted the word-by-word elicited MEG signal in a widespread cortical network, irrespective of sentential context. In contrast, index (ordinal word position) was more strongly encoded in sentence words, in left front-temporal areas. This confirms that frequency influences word processing independently of predictability, and that contextual constraints affect word-by-word brain responses. With a conservative multiple comparisons correction, only the interaction between lexical frequency and surprisal survived, in anterior temporal and frontal cortex, and not between lexical frequency and entropy, nor between lexical frequency and index. However, interestingly, the uncorrected index*frequency interaction revealed an effect in left frontal and temporal cortex that reversed in time and space for intact compared to scrambled sentences. Finally, we provide evidence to suggest that, in sentences, lexical frequency and predictability may independently influence early (<150ms) and late stages of word processing, but also interact during late stages of word processing (>150-250ms), thus helping to converge previous contradictory eye-tracking and electrophysiological literature. Current neuro-cognitive models of reading would benefit from accounting for these differing effects of lexical frequency and predictability on different stages of word processing.

## 1. Introduction

When reading a text, the reader’s brain is capable of rapidly extracting meaning from the structured sequence of individual words. In order to achieve its remarkable efficiency in processing, the brain network for language not only extracts, and actively uses, lexical properties of the individual words, but is also greatly influenced by the context in which those words occur. On the one hand, for instance, words that maintain a highly frequent occurrence in day-to-day language use are processed faster and with less effort than words that occur less frequently (Calvo & Meseguer, 2002; Inhoff & Rayner, 1986; Rayner & Duffy, 1986; Rubenstein, Garfield, & Millikan, 1970). On the other hand, as a linguistic expression unfolds, the previously read input provides the brain with a constraining semantic and syntactic context, which may allow for predictions to be made about the upcoming word. This results in measurable effects at fast timescales, in response times (Staub, Grant, Astheimer, & Cohen, 2015), and in both electrophysiological (Van Petten & Kutas, 1990) and eye movement signals (Calvo & Meseguer, 2002).

Typical adult readers effortlessly process an average of 238 words per minute (Brysbaert, 2019), fixating on each word for an average of only 235ms (Rayner, 1986). The brain’s rapid word processing has been shown to be facilitated when the word frequently occurs within a given language (i.e. has a high *lexical frequency*). Compared to low frequency words, high frequency words are fixated for shorter durations during reading (Calvo & Meseguer, 2002; Inhoff & Rayner, 1986; Rayner & Duffy, 1986), are responded to faster in lexical decision tasks (Rubenstein et al., 1970), and produce smaller electrophysiological (Smith & Halgren, 1987; Van Petten & Kutas, 1990) and hemodynamic responses (Chee, Hon, Caplan, Lee, & Goh, 2002). Although the specific temporal and spatial dynamics of electrophysiological frequency effects may be sensitive to task context (Chen, Davis, Pulvermüller, & Hauk, 2015; Strijkers, Bertrand, & Grainger, 2015), overall, it seems that processing of high frequency words is less effortful than low frequency words.

The prediction of upcoming sentential content is another mechanism that seems to facilitate the remarkable speed of sentence reading. There is now ample evidence that one is able to predict upcoming linguistic input, although whether this is to the level of semantics, syntactic content or the word form is still debated (Pickering & Gambi, 2018). Regardless of the level at which prediction takes place, highly predictable words seem to be processed faster than unpredictable words, reflected in shorter fixation durations (Calvo & Meseguer, 2002; Rayner & Well, 1996) and smaller N400 responses (Van Petten & Kutas, 1990). The N400 is an electrophysiological marker of semantic processing, which occurs between 200-600ms at a centro-parietal topography, and is thought to reflect either the integration and unification of semantic information (Hagoort, Baggio, & Willems, 2009; Kutas & Federmeier, 2011) or conceptual (or possibly lexical) pre-activation (Lau & Namyst, 2019; Lau, Phillips, & Poeppel, 2008). A larger N400 response is observed when the integration of semantic information is more difficult, or in the absence of conceptual/lexical pre-activation, for example when the word is less predictable.

There is increasing agreement that there are two mechanisms through which prediction can take place. Firstly, through a fast, effortless and automatic mechanism, in which activity spreads to associated features, or, secondly, through a higher level mechanism, in which world knowledge and the surrounding context are combined to form predictions (Huettig, 2015; Pickering & Gambi, 2018). Lexical frequency could therefore influence the automatic, bottom-up prediction mechanism, where activation thresholds are lower for high compared to low frequency words. In contrast, effects of the semantic and syntactic constraints, provided by the context that a word is presented in, may reflect a prediction mechanism that relies on the top-down flow of information from strong priors. For example, as semantic context increases as the sentence unfolds a stronger foundation on which to base predictions is provided. In this study, we follow earlier approaches in using ordinal word position in a sentence (or *index*) to roughly quantify context. Indeed, the N400 has been shown to decrease with both increased lexical frequency and increased index (Dambacher, Kliegl, Hofmann, & Jacobs, 2006; Payne, Lee, & Federmeier, 2015; Van Petten & Kutas, 1990), which suggests that word integration becomes easier as each of these factors increase.

A recurring finding in the literature is that there is an interaction between effects of increased predictability and lexical frequency on the N400, where the effect of word frequency on the N400 amplitude during word processing is greatly diminished or disappears with increased context or predictability (Alday, Schlesewsky, & Bornkessel-Schlesewsky, 2017; Dambacher et al., 2006; Payne et al., 2015; Sereno, Hand, Shahid, Mackenzie, & Leuthold, 2019; Van Petten & Kutas, 1990). Similar interactions have also been observed at earlier time windows (Dambacher et al., 2012; Sereno, Brewer, & O’Donnell, 2003; Sereno et al., 2019) and with functional near-infrared spectroscopy (fNIRS; Hofmann et al., 2014). In an MEG study, Fruchter, Linzen, Westerlund, and Marantz (2015) additionally found word frequency and predictability to interact in the left MTG, in time windows both preceding and succeeding the predictable word onset. Overall, these findings demonstrate that the interaction between lexical frequency and increased context is a robust and well replicated finding, which reflects both the reduced influence of lexical frequency on word processing with increased context, as well as a greater benefit of predictability for processing low compared to high frequency words.

The reduced effect of lexical frequency on word processing with increased context has lead authors to conclude that lexical frequency merely reflects a bottom-up, baseline level of expectation that is soon overridden with top-down information in the presence of context (Kretzschmar, Schlesewsky, & Staub, 2015). However, there is a well-documented discrepancy between the aforementioned electrophysiological literature and the eye-tracking literature as to whether frequency and predictability indeed have an interactive effect on word processing, or whether effects are additive (Kretzschmar et al., 2015). In contrast to the findings of the N400 literature, recording participants’ eye gaze during reading has consistently demonstrated an additive effect of lexical frequency and predictability on fixation durations. Fixation durations are longer for highly predictable low frequency words than highly predictable high frequency words, and again longer for unpredictable low frequency words (Kennedy, Pynte, Murray, & Paul, 2013; Kretzschmar et al., 2015; Staub, 2015; Staub & Benatar, 2013). One explanation for these contradictory findings is that lexical frequency and prediction have separate additive effects during early processing stages (Sereno et al., 2019; Staub & Goddard, 2019), for example during sublexical orthographic processing, morphological decomposition or lexical retrieval, but that frequency effects are not present with increased context during later semantic processing and integration.

### 1.1. The current work

Considering the aforementioned ambiguity in the theoretical understanding of how lexical frequency influences subsequent processing, specifically in the light of additional context-based predictability, the current work performed a novel analysis on an existing dataset, with the aim to dissociate lexical frequency effects from predictability effects. Although previous work has sought to define *when* frequency and predictability interact, less attention has been invested into examining the spatiotemporal dynamics of this interaction (although, see the exploratory analysis in Fruchter et al., 2015 for an exception). We aimed to determine at which time points and in which locations lexical frequency and predictability independently influence word processing, and at which points they interact, thereby providing valuable information for models of word reading. Staub and Goddard (2019) recently highlighted that current models of word reading, such as the E-Z reader (Reichle, Rayner, & Pollatsek, 2003) and SWIFT (Engbert, Nuthmann, Richter, & Kliegl, 2005), do not yet completely account for effects of predictability and invalid previews on fixation durations. Considering the complex effects lexical attributes have on the neural processing of language, a comprehensive account of word reading could benefit from improving upon both the temporal and spatial resolution of previous work.

Specifically, we used the Mother of all Unification Studies (MOUS; Schoffelen et al., 2019), a large sample size open-access dataset of 102 participants in which magnetoencephalography (MEG) was recorded while they read intact sentences and scrambled sentences. Improving upon previous electroencephalography (EEG), functional magnetic resonance imaging (fMRI) and fNIRS research, MEG provides both the temporal and spatial resolution to detect subtle and fine-grained differences in the extent that lexical frequency and predictability are encoded in the MEG signal after word-onset, which could have previously been lost by averaging over time and space. Distinct from most previous work, with respect to the analysis, we exploited the word-by-word variability in the MEG signal, which is often lost through averaging across words of the same experimental condition. Specifically, we used multiset canonical correlation analysis (MCCA) to boost the stimulus-specific signal (Arana, Marquand, Hultén, Hagoort, & Schoffelen, 2020), and performed detailed cross-validated single-trial encoding model analysis, using regression models that quantified the degree to which lexical frequency and various measures of predictability are encoded in the MEG signal.

To investigate the extent that context influences effects of lexical frequency on word processing, we first compared sentences and scrambled sentences as to the amount of variance in the ongoing brain signal explained by lexical frequency. The *scrambled* sentences were created by randomly shuffling the order of the words in the *intact* sentences, and therefore matched the intact sentences word-for-word, differing only in the order that words were presented in. This meant that the two conditions (intact/scrambled) differed only in the presence/absence, respectively, of the build-up of a rich sentence context. Although some degree of sparse combinatorial processing may have been possible at the semantic level in the scrambled sentences, the ability to derive a coherent sentence level context and produce top-down driven predictions was possible only in the sentences. In addition to the level of sentential context provided by the presence/absence of syntax, we approximately quantified context with the ordinal word position in the sentence (index), consistent with previous approaches (Dambacher et al., 2006; Payne et al., 2015; Van Petten & Kutas, 1990). Index captures the incremental build-up of the entire sentence context. Moreover, as context increases with increased word position, predictability is expected to increase with increased context (for a similar argument, see Levy, 2008; Schuster, Hawelka, Himmelstoss, Richlan, & Hutzler, 2020). Thus, effects of index were expected to differ in intact compared to scrambled sentences. Whereas index provided a correlate of predictability that encompassed the entire sentence context, surprisal and entropy were used to provide measures of local predictability (acquired from a trained tri-gram model). Specifically, surprisal quantifies how unexpected the current word is, and entropy represents the uncertainty of the upcoming word. Effects of surprisal and entropy were compared across intact and scrambled sentence conditions, in order to identify effects related to higher level predictive processes, which were only possible in the sentence condition. We investigated the interaction between lexical frequency and each variable quantifying different degrees of predictability (index, surprisal and entropy). Lexical frequency (rather than lemma frequency) was chosen to quantify word frequency effects, in order to remain consistent with most previous reports (Alday et al., 2017; Dambacher et al., 2006; Payne et al., 2015; Sereno et al., 2019; Van Petten & Kutas, 1990). As effects of lexical frequency and predictability on the electrophysiological response have been shown to interact with word length (Penolazzi, Hauk, & Pulvermuller, 2007), word length was added as a control predictor to all models. Due to fundamental differences in the properties of content words (nouns, adjectives, verbs) and function words (determiners, prepositions, pronouns, conjunctions), for example in their frequency, length and semantic richness, they were analysed separately (see Matchin, Brodbeck, Hammerly, & Lau, 2019 for a similar approach). Only content words were included in the analysis here.

Although Fruchter et al. (2015) previously studied the spatiotemporal effects of a similar interaction using MEG, our study differed from theirs in a number of ways, providing additional contributions to the field. Firstly, in contrast to Fruchter et al. (2015), our stimuli were not designed to be highly predictable, and were not limited to measuring the response to adjective-noun pairs such as “stainless steel”, selected based on co-occurrence statistics. We therefore investigated the spatiotemporal dynamics of the interaction with a richer stimulus set, which is arguably closer to the linguistic content one would read in everyday situations, where sentences are not always highly predictable, and also depend upon integrating world knowledge. The prediction of frequently co-occurring words would arguably depend on different processing mechanisms (e.g. priming) compared to forming predictions based on the build-up of context constraints (Huettig, 2015; Pickering & Gambi, 2018). Secondly, we investigated the effect of the interaction over time and space, rather than averaging over time windows or using single regions-of-interests (ROIs). Although Fruchter et al. (2015) also presented the spatiotemporal dynamics of the interaction, this was in an exploratory analysis that requires replication. Their primary analyses averaged over longer time windows and were restricted to several ROIs. Furthermore, in their exploratory spatiotemporal analysis, the authors averaged over 100ms time windows. We here provide finer grained information about the spatiotemporal dynamics of the interaction between lexical frequency and context. Finally, we investigated whether such effects were observable on the level of word-by-word processing during sentence reading, without averaging over trials, by quantifying the improvement of MEG signal prediction in a comparative cross-validated model scheme.

## 2. Methods

### 2.1. Participants

Participants were 99 right-handed native Dutch speakers (age range 18-33 years; mean age = 22; 50 males) from a subset of 102 participants who completed a reading paradigm in the open-access MOUS dataset (Mother of all Unification Studies; Schoffelen et al., 2019). Three participants were excluded from analyses, due to technical issues during data acquisition making them unsuitable for the current analysis pipeline. All participants were right handed, had normal or corrected to normal vision, and reported no history of neurological, developmental or language impairments. All participants provided written informed consent and the study was approved by the local ethics committee, and complied with the declaration of Helsinki.

### 2.2. Sentence stimuli

The total stimulus set consisted of 360 Dutch sentences (9-15 words in length), which are described in detail in Schoffelen et al. (2019). Each participant read a selection of 240 sentences (2/3 of the entire stimulus set), where 50% were presented as intact sentences and 50% were presented as scrambled sentences. Specifically, three pairs of selections, referred to as *scenario pairs*, were created, such that the stimuli that occurred as normal sentences in one scenario from a pair were presented in a scrambled fashion in the other scenario from that pair, and vice versa. Sentences were scrambled so that no more than three words in a scrambled sentence made up a coherent phrase. No participant read both the intact and scrambled version of a sentence. Consequently of this design was that the collection of words that subjects read was exactly counterbalanced across intact and scrambled sentence conditions, both across all participants and within the three sets of scenario pairs.

### 2.3. Lexical characteristics

Lexical characteristics of frequency, index, surprisal, entropy and length (i.e. number of characters) were obtained for each word in the sentence, to enter as predictors into regression models. *Lexical frequency* was defined as the frequencies of words occurring in the NLCOW2012 corpus (Schäfer & Bildhauer, 2012) and were log10 transformed. The NLCOW2012 database is comprised of over 10 million Dutch sentences (71761868 words), and was also used to obtain estimates of surprisal and entropy (see below). *Index* was defined as the ordinal position of the word in the intact/scrambled sentence. Each word’s *surprisal* value was acquired from a trained tri-gram model, using WOPR (Van Den Bosch & Berck, 2009), trained on the NLCOW2012 corpus. Surprisal was computed as the conditional probability of observing a word given the previous two words in the sentence. Formally, it was computed as:

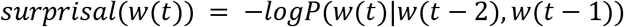

High surprisal values therefore signify low lexical predictability. *Entropy* was acquired from the same trained tri-gram model. Entropy reflects the probability distribution of possible continuations, given the constraints of the previous words. High entropy values signify a high number of possible continuations, i.e. low predictability of the upcoming word. Formally, it is defined as:

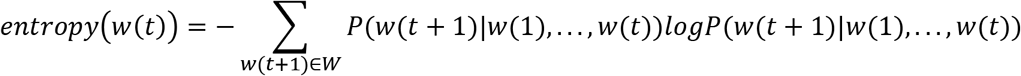

Using a trained tri-gram model here, the entropy at word w(t+1) reflects the summation across all possible endings given w(t) and w(t-1).

All metrics were also computed for the first two words in a sentence. The statistical language model allowed for estimates of sentence onset words (and also the second word in the sequence), since sentences were prepended by special tokens, which allowed the first sentence word to be treated as a valid trigram.

The distribution of the estimated surprisal values for both scrambled and intact sentences are presented in Fig 1. Here it can be seen that model-based surprisal and entropy are higher for scrambled than intact sentences. Although there were likely many trigrams in the current stimuli that were not present in the corpus on which the language model was trained, particularly in the scrambled sentence condition, N-gram based statistical language models account for this by estimating the conditional probabilities using a technique called smoothing (or discounting), returning non-zero probabilities for words, even if corresponding trigrams did not occur in the training set. In such cases, the returned conditional probabilities will be more closely related to the (unconditional) lexical frequency of the word. Fig 1 additionally highlights that measures of lexical frequency and surprisal, and lexical frequency and length were highly correlated. This is unsurprising, as both lexical frequency and surprisal were calculated from the frequency of occurrences in a corpus, either of the word itself, or the word given the prior two words. Such high correlations were not a concern for the current analysis, in which we used a model comparison procedure to quantify the additional variance explained by a model including the independent variable compared to a reduced model that did not contain the independent variable. A detailed explanation of the model comparison procedure can be found in section 2.7.

**Figure 1.**
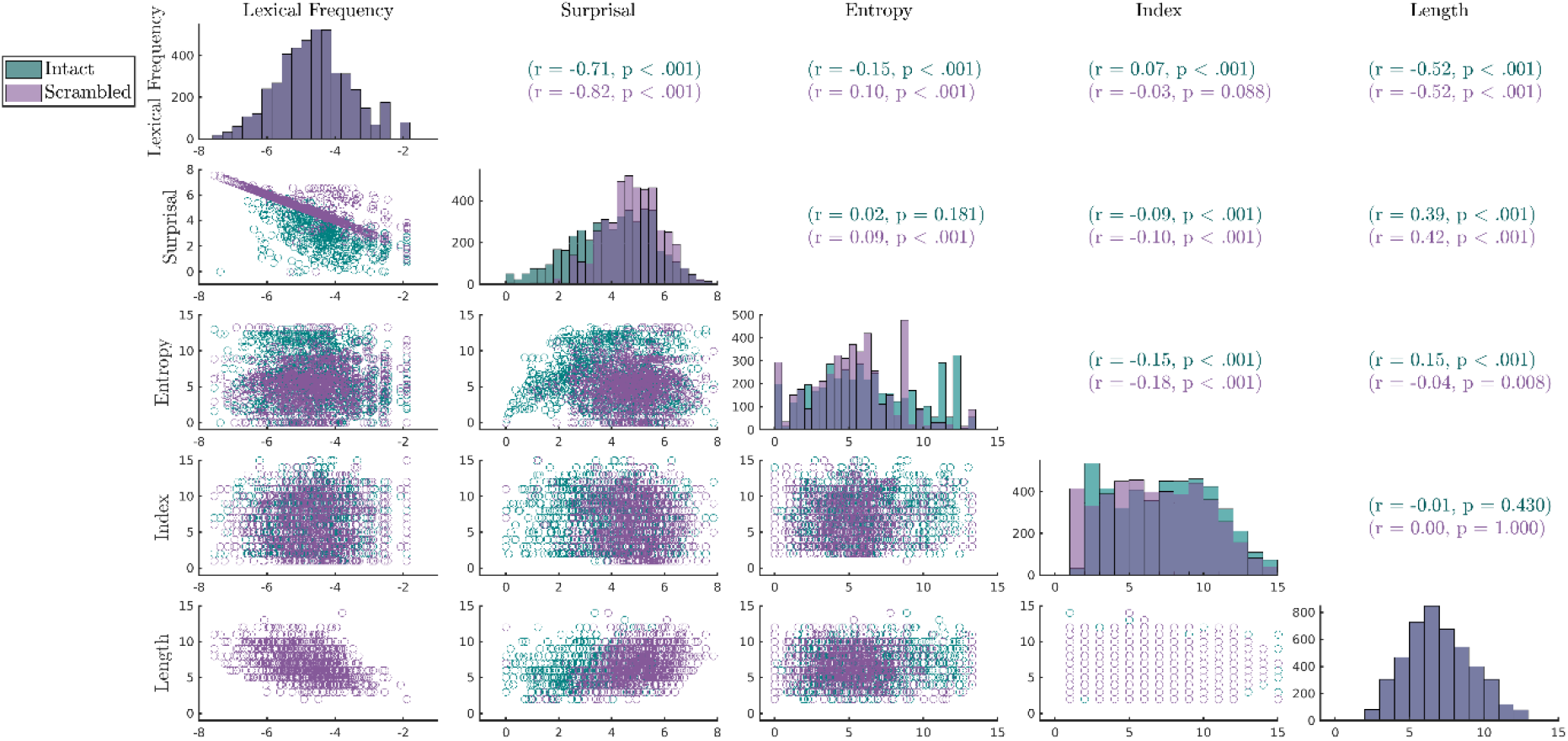
Correlation matrix for predictor variables lexical frequency (log10-transformed), surprisal (log10- transformed), entropy, index and word length (respectively) for the content words. Scatterplots between corresponding pairs of predictors are presented in the lower off-diagonal. Pearson’s correlation coefficients and corresponding p values are presented on the upper off-diagonal. Histograms present the distribution of each predictor variable on the diagonal.

### 2.4. Experimental procedure

Sentence stimuli were presented in a random order in alternating intact and scrambled sentence blocks. There were 48 blocks in total, each containing five intact sentences or five scrambled sentences. The starting block condition (intact/scrambled) was randomised across participants. At the beginning of each block the block type was presented on the screen for 1500ms. Trials (intact/scrambled sentences) were separated with a 1200-2200 inter-trial interval, during which a blank screen was presented followed by a fixation cross. Stimuli were presented word-by-word, with an inter-stimulus (word) interval of 300ms. To avoid the entrainment of neural oscillations to a rhythmic onset of visual stimuli, and to better match the pace of the equivalent spoken stimuli (Schoffelen et al., 2019), the presentation duration of each word was adjusted by the word duration when spoken (visual presentation duration = 300-1400ms, mean = 351ms). The calculation of single word durations has been described elsewhere (Lam, Schoffelen, Udden, Hulten, & Hagoort, 2016; Schoffelen et al., 2019). To reiterate, for each intact/scrambled sentence, the duration of a single word was a function of four factors: (i) the duration of the spoken version of the intact/scrambled sentence in the matching auditory stimuli from Schoffelen et al. (2019) (audiodur), (ii) the total number of words in the sentence (nwords), (iii) the number of letters per word (nletters), and (iv) the total number of letters in the sentence (sumnletters). Single word duration was computed as: (nletters/sumnletters)*(audiodur+2000–150*nwords)

The minimum presentation duration for short words was limited to 300ms, regardless of the outcome of the above formula. As the presentation rate of stimuli was partially determined by the refresh rate of the projector (60Hz), the actual presentation duration of words increased by 0-33ms from the value provided by the above formula.

Participants were instructed to read the sentences. On 20% of trials participants answered a yes/no comprehension question to ensure they were engaged in the task. The positions of the comprehension questions relative to the stimuli were random. In intact sentence blocks, 50% of questions asked about the content of the sentence (e.g. “Did grandma give a cookie to the girl?”). Questions in the scrambled sentence blocks, and the remaining 50% of questions in the intact sentence blocks, asked about the presence of a content word (e.g. “Was the word *grandma* mentioned?”). Participants responded to the questions by pressing a button with their left index/middle finger to answer yes/no, respectively.

Stimuli were presented with Presentation software (Version 16.0, Neurobehavioral Systems, Inc) and back-projected with an LCD projector at a refresh rate of 60Hz. Words were presented in the centre of the screen in a black mono-spaced font (visual angle of 4 degrees) on a grey background. Before beginning the main experiment, participants completed practice trials to familiarise themselves with the procedure.

### 2.5. MEG acquisition

Participants were seated in a magnetically shielded room, while MEG was recorded with a 275 axial gradiometer CTF system, at a sampling rate of 1200 Hz and with a 300 Hz analog low pass filter. Prior to the recording, the participant’s head shape was digitised with a Polhemus 3D-Space Fast-track digitiser. Digitised head shapes and fiducial points were later used to coregister subject-specific anatomical MRIs with the MEG sensor space. The position of the participants’ head (relative to the MEG sensors) was monitored online throughout the recording via three head-localiser coils, placed on the nasion and left and right pre-auricular points.

### 2.6. MRI acquisition

MRIs were recorded with a Siemens Trio 3T MRI scanner with a 32-channel head coil. A T1-weighted magnetisation-prepared rapid acquisition gradient echo pulse sequence was used to obtain structural MRIs (volume TR = 2300ms; TE = 3.03ms; 8° flip angle; 1 slab; slice matrix size = 256 × 256; slice thickness = 1mm; field of view = 256mm; isotropic voxel size = 1.0 × 1.0 × 1.0mm). A vitamin E capsule was placed behind the right ear as a fiducial marker to visually identify left/right.

### 2.7. Data analysis

#### Pre-processing

Data were band pass filtered between 0.5-20Hz and epoched time-locked to sentence onset. Segments of data that contained eye blinks, squid jumps and muscle artifacts were replaced with “Not a Number” (NaN) in order to preserve the original sentence onset related timing information. Data were downsampled to 120Hz.

#### Source Reconstruction

Single shell head models describing the inside of the skull were constructed from individual MRIs, which were used to create forward models according to (Nolte, 2003). Single trial covariance matrices were computed between sensor pairs. Sources were reconstructed using linearly constrained minimum variance (LCMV; Van Veen, van Drongelen, Yuchtman, & Suzuki, 1997) beamforming to obtain time courses of source activity at 8196 dipole locations. Data were parcellated using an anatomical atlas-based parcellation, consisting of 382 parcels (Schoffelen et al., 2017). For each parcel, principal component analysis was performed on the dipole time series belonging to a given parcel, and the top five components that explained the most variance in the parcel-specific signal were selected for further analysis.

#### Spatiotemporal Alignment

To boost the stimulus specific signal, and reduce intersubject variability, data were spatiotemporally aligned across subjects using multiset canonical correlation analysis (MCCA; Arana et al., 2020; de Cheveigné et al., 2019). MCCA was used to find linear combinations of the 65 parcel time courses (canonical components) that maximised the correlation between all subject pairs, while they were presented with exactly the same words, thereby increasing the similarities between the participants’ signals in response to those words.

MCCA is a generalization of canonical correlation analysis (CCA), and aims to find linear combinations for multivariate observations in order to maximize the correlation between the combined time series. Here, each member of the set of multivariate observations consisted of a representation of a parcel-specific signal for a given subject. Linear combinations of these observations were estimated, which resulted in a single canonical component per subject such that the correlation across subjects was maximised. The linear weights were estimated with a generalized eigenvalue decomposition using two covariance matrices, consisting of the full covariance matrix of all subjects’ multivariate observations, and of a block-diagonal covariance matrix, containing only the within subject covariances of the multivariate observations. As mentioned, our aim was to boost the stimulus-specific brain signals, specifically accounting for some spatial and temporal variability across subjects. Hence, for each subject the input to MCCA decomposition consisted of a set of time-shifted time series, where the parcel’s 5 dominant principal components were shifted in time from -50-50ms in steps of single samples, resulting in 65 time series per word per parcel and subject (i.e. 5 principal components × 13 time shifts). MCCA was performed separately for each pair of scenarios, which were fully matched in terms of the stimulus material that was used to derive the sentences and the word-lists (i.e. the subjects read exactly the same overall collection of individual words), based on combining data from sets of 32-34 subjects. Next, the time series of the scrambled sentence trials were unscrambled such that the word order and onset times exactly matched the corresponding intact sentence’s word order and onset times. This resulted in 240 trials that were exactly matched across time in terms of the individual words presented. These trials were entered into a five-fold cross validated MCCA procedure (Arana et al., 2020). To this end, we partitioned the data into 5 test folds of 48 trials each, and for each of the folds used the 192 remaining trials as a training set to estimate the MCCA weights. These weights were subsequently applied to the test fold data to obtain the subject-specific canonical components. The cross validation was applied in order to avoid overfitting. To summarise, MCCA was used to find linear combinations of the 65 parcel time courses (canonical components) that maximised the correlation between all subject pairs, while they were presented with exactly the same words, thereby increasing the similarities between the participants’ signals in response to those words.

#### Encoding Models

Next, we fitted encoding models to the data, using five-fold cross-validated ridge regression. To this end, the subject-specific canonical components were re-epoched time-locked to word onset, selecting only content words (nouns, adjectives, and verbs). The content words made up 55% of all the words in the stimulus set, which resulted in an average of 763 (range: 755-774) words per scenario and main condition (intact versus scrambled sentences). The absolute number of analysed words per sentence varied as a function of sentence length. For the re-epoched data, subject-specific encoding models were estimated for each time point and parcel-of-interest, separately for intact and scrambled sentence words. A ridge regression model is similar to a multiple regression model with a regularised design covariance matrix. The optimal regularisation parameter was estimated using nested cross-validation, and selected from a range of lambda values (0.002, 0.005, 0.010, 0.020, 0.050, 0.100, 0.200, 0.500, 1.000, 2.000 and 5.000) for each model. The regularisation parameter applies a penalty to the model to avoid overfitting on the training data. A lambda value of 0 would result in no regularisation being applied, whereas selecting a lambda value that is too high would result in under-fitting the model. The model derived from a “training” portion of the data was evaluated on its performance to predict a portion of unseen “test” data.

In order to separate the unique variance explained by each variable of interest from that explained by all other variables, we applied a model comparison scheme. The model comparison procedure quantified the extent to which a model including a predictor of interest explained variance in the MEG signal, above and beyond a reduced model that did not include the given predictor. To this end we computed the coefficient of determination:

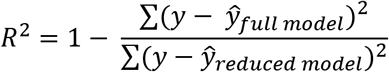

Where the numerator and denominator in the right side of the equation were computed as the sum-of-squares of the difference between the data and the modelled test data, for the full and reduced models, respectively.

To test the contribution of individual predictors we used a full model that included, beyond a constant and word length, the following predictors of interest: lexical frequency (log transformed), surprisal, entropy and index. To test the interaction between lexical frequency and context – as quantified with index (similar to Alday et al., 2017; Payne et al., 2015; Van Petten & Kutas, 1990) – we used a full model that included only, beyond a constant, the individual predictors of lexical frequency (log transformed), index, length, and the interaction term, which was computed as an element wise product: lexical frequency (log10 transformed) × index. Similarly, we tested the interaction between lexical frequency and surprsial, and lexical frequency and entropy, where the full model included, beyond the constant, the individual predictors of lexical frequency (log transformed), surprisal (log transformed)/entropy, length, and the interaction term (lexical frequency × surprisal (log transformed)/lexical frequency × entropy). Epochs (content words) were divided into five equal folds to avoid overfitting, and to allow for the generalisation across items. For each fold of the cross-validation procedure, the model was estimated using data from the four other folds, and tested on the remaining data.

In order to be able to statistically compare the models for the individual intact and scrambled sentence conditions, that is to obtain an estimate of a possible bias in the coefficient of determination under the null hypothesis, we used a permutation approach, as follows: For each model, the design matrix was randomly permuted 50 times and, for each permutation, an additional model was trained and tested with the permuted variables, thereby removing any true association between the predictors and the data.

#### Statistical Analysis

We statistically evaluated the individual predictors in a selection of regions-of-interest (ROI), consisting of 184 parcels (92 left hemisphere parcels with their right hemisphere counterparts). This selection consisted of cortical regions that have consistently been described to be a part of a language network (Catani et al., 2007; Friederici, 2009; Glasser & Rilling, 2008; Schoffelen et al., 2017) or to be involved in the processing of semantic relationships (Bunge, Helskog, & Wendelken, 2009; Frankland & Greene, 2020; Knowlton, Morrison, Hummel, & Holyoak, 2012; Ramnani & Owen, 2004). We further investigated the interaction between lexical frequency and index based on the resulting map including only the 33 parcels that significantly encoded index or lexical frequency. The interaction between lexical frequency and surprisal, and lexical frequency and entropy were investigate in the same 33 parcels, facilitating comparison across results.

We used non-parametric permutation statistics, using the dependent samples T-statistic across subjects as a test statistic. We evaluated the individual coefficients of determination against the corresponding average of their 50 random permutation counterparts (see Encoding Models section 2.7), using an alpha-level of 0.05 for inference. The intact and scrambled sentence conditions were compared with each other using a two-sided test (which involves evaluating the test statistic against two randomisation distributions, using an alpha level of 0.025 for each of these randomisation distributions) for inference. For all comparisons, multiple comparisons (across time and space) were accounted for by using a max-statistic distribution from 5000 permutations.

Note that we compared intact and sentence conditions only on the difference in the interaction between lexical frequency and index, and not in the interaction between lexical frequency and surprisal, nor lexical frequency and entropy. Index is well-controlled across intact/scrambled sentence conditions, in that it is well matched across both intact and scrambled sentences, and does not correlate with lexical frequency. In contrast, the distribution of surprisal and entropy both differ across intact and scrambled sentences, with higher surprisal and entropy values in scrambled compared to intact sentences (see Fig 1). Any observed difference between intact and scrambled sentences in the variance explained by the interaction between lexical frequency and surprisal/entropy could, therefore, be down to their different distributions of surprisal/entropy values.

## 3. Results

All participants achieved over 60% accuracy on the comprehension questions (mean = 81.19%; sd = 6.61%), confirming they were attending to the stimuli. No further analysis was conducted on the comprehension questions.

### 3.1. Spatiotemporal Alignment

Fig 2 shows the effect of the alignment procedure, presenting the time-resolved intersubject correlation (Fisher Z-transformed correlation coefficient) after spatiotemporal alignment (solid green line), spatial alignment (dashed green line), temporal alignment (dotted red line) and no alignment (dashed purple line), for two example parcels (sub-regions of BA22 and BA44). Fig 2 illustrates that spatiotemporal alignment increased the intersubject correlation, more so than temporal alignment alone or spatial alignment alone. The intersubject correlation peaked at around 400ms (300-500ms), a time period in which electrophysiological brain signal is typically found to be influenced by the semantic characteristics of a word (N400/M400; Kutas & Federmeier, 2011). Spatiotemporal alignment thereby seems to have boosted the stimulus specific signal in the data.

**Figure 2.**
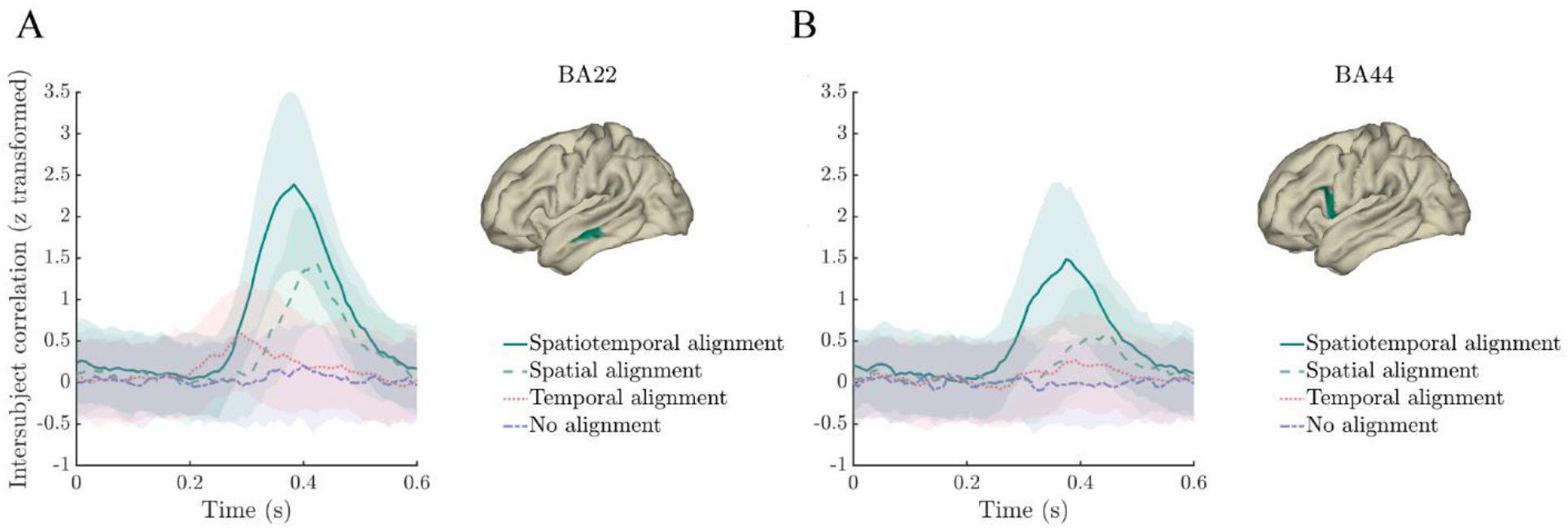
MCCA boosts intersubject consistency of single word responses. Time courses of Z-transformed intersubject correlations after spatiotemporal alignment (solid green line), spatial alignment (dashed green line), temporal alignment (dotted red line) and no alignment (dashed purple line) in middle temporal gyrus (parcel in Brodmann Area (BA) 22; panel A) and inferior frontal gyrus (parcel in BA44; panel B). Shaded ribbons represent the interquartile range.

### 3.2. Encoding Models

For each measure of interest, our model comparison scheme quantified the extent that each regressor explained word-specific variance in the MEG signal, beyond the variance explained by all other regressors (see Methods section 2.7). Similarly, we quantified the variance explained by the index × lexical frequency interaction, surprisal × lexical frequency interaction and entropy × lexical frequency interaction, beyond that explained by the main effects of lexical frequency and index/surprisal/entropy (respectively). The model comparisons were statistically evaluated separately for the intact (Figs 3-9 panel A) and scrambled (Figs 3-7 panel B) sentences against a permutation derived baseline, as well as compared against each other (Figs 3-7 panel C).

**Figure 3.**
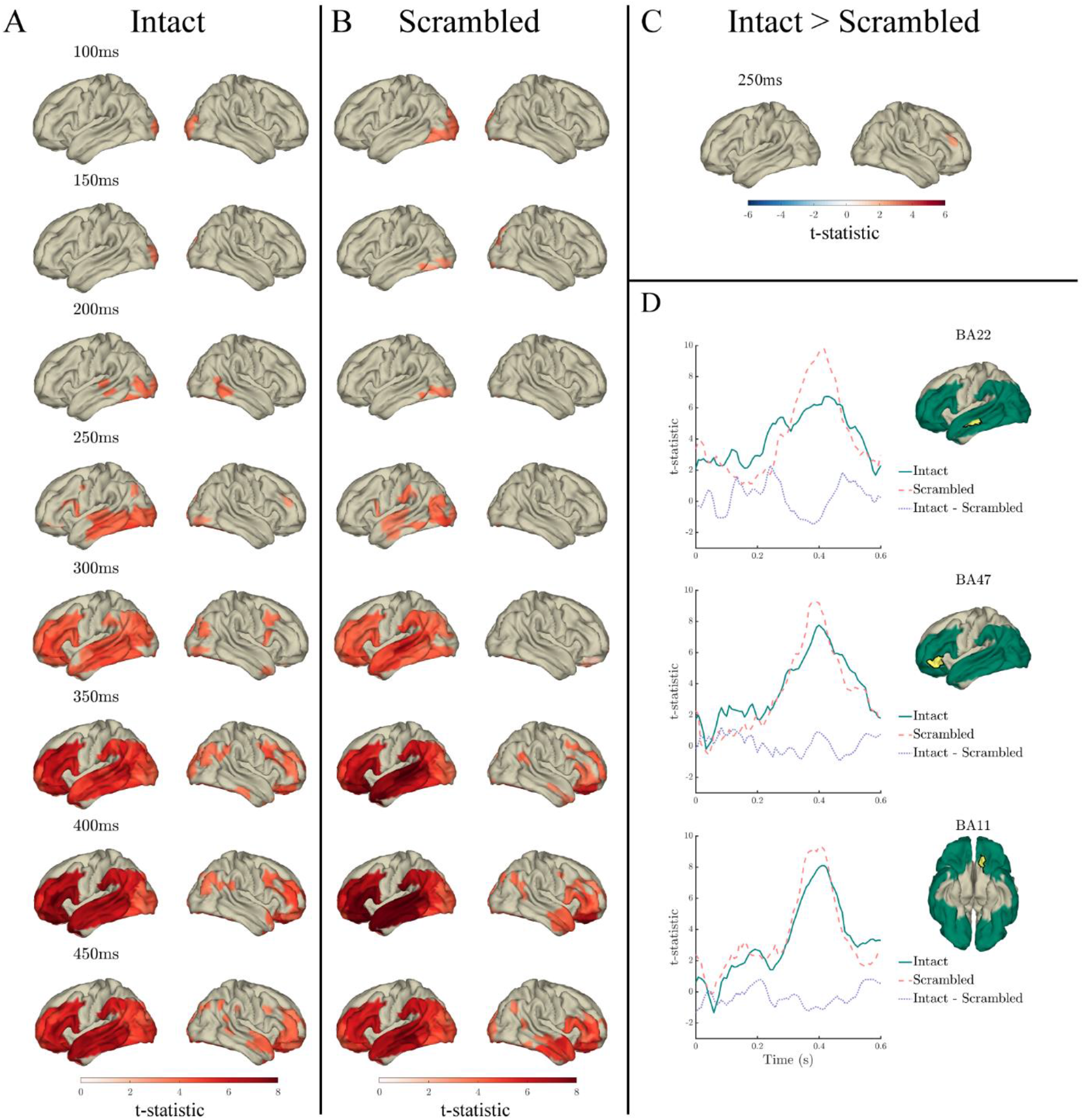
Effects of lexical frequency in the response to content words: Surface plots of T-statistics (averaged over 50ms time windows centred at the indicated latencies, for visualisation) quantifying the difference in variance explained by lexical frequency (log10 transformed), beyond that explained by index, surprisal, entropy and length, in intact sentence compared to random permutation models (panel A; *p*<.05 one-sided, corrected), scrambled sentence compared to random permutation models (panel B; *p*<.05 one-sided, corrected), and intact compared to scrambled sentence models (panel C; *p*<.05 two-sided, corrected). Parcels for which no time point was significant during the 50ms time bin are masked. Panel D: Time courses of T-statistics for intact (solid green line) and scrambled (dashed red line) sentence models compared to random permutation models, and intact compared to scrambled sentence models (dotted purple line) for subparcels of BA22, BA47 and BA11 (highlighted in yellow on adjacent surface plots). ROIs entered into statistical analyses are illustrated as green shaded area on surface plots.

**Figure 4.**
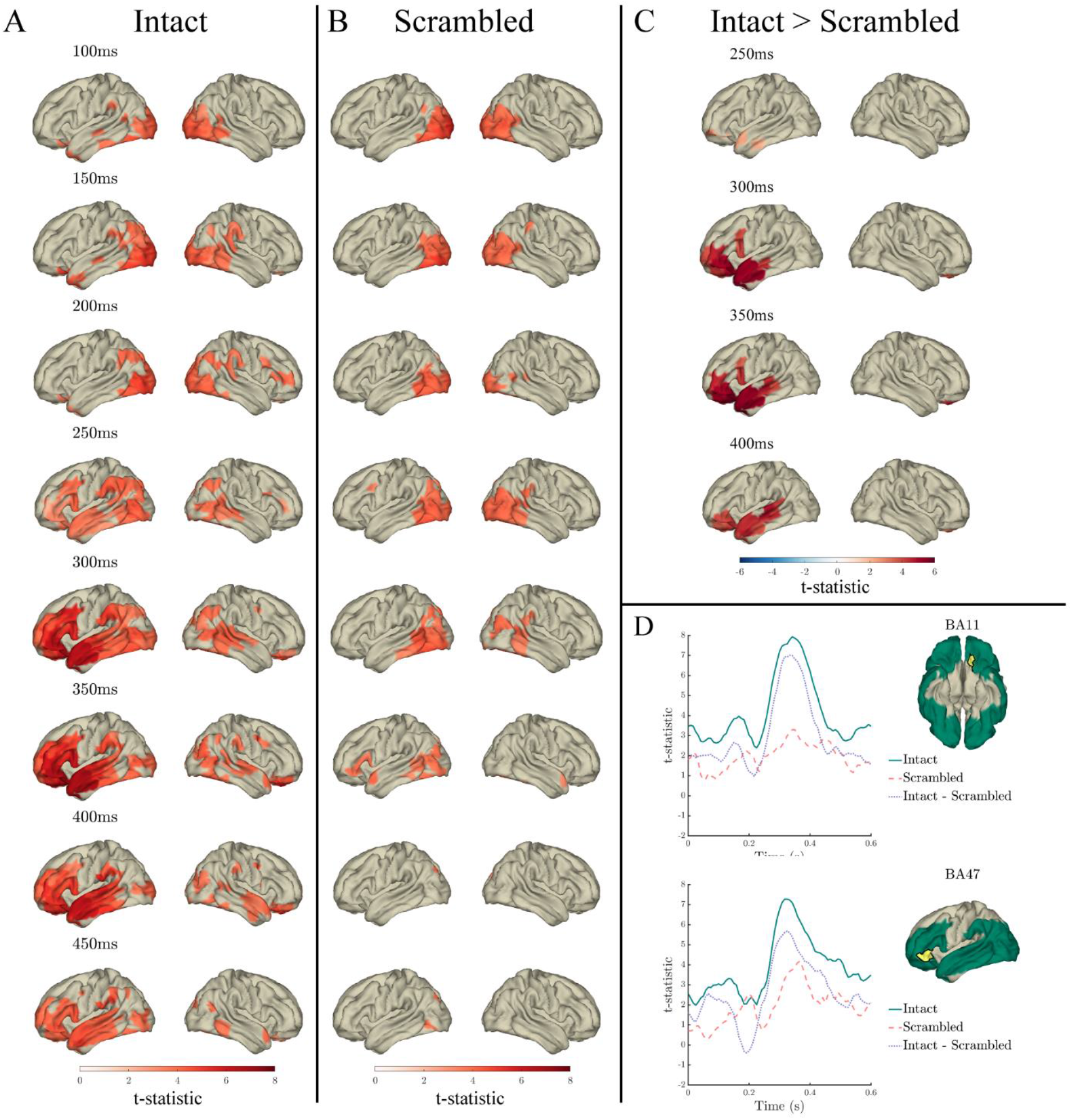
Effects of index in the response to content words: Surface plots of T-statistics (averaged over 50ms time windows centred at the indicated latencies, for visualisation) quantifying the difference in variance explained by index, beyond that explained by lexical frequency (log10 transformed), surprisal, entropy and length, in intact sentence compared to random permutation models (panel A; *p*<.05 one-sided, corrected), scrambled sentence compared to random permutation models (panel B; *p*<.05 one-sided, corrected), and intact compared to scrambled sentence models (panel C; *p*<.05 two-sided, corrected). Parcels for which no time point was significant during the 50ms time bin are masked. Panel D: Time courses of T-statistics for intact (solid green line) and scrambled (dashed red line) sentence models compared to random permutation models, and intact compared to scrambled sentence models (dotted purple line) for subparcels of BA11 and BA47 (highlighted in yellow on adjacent surface plots). ROIs entered into statistical analyses are illustrated as green shaded area on surface plots.

**Figure 5.**
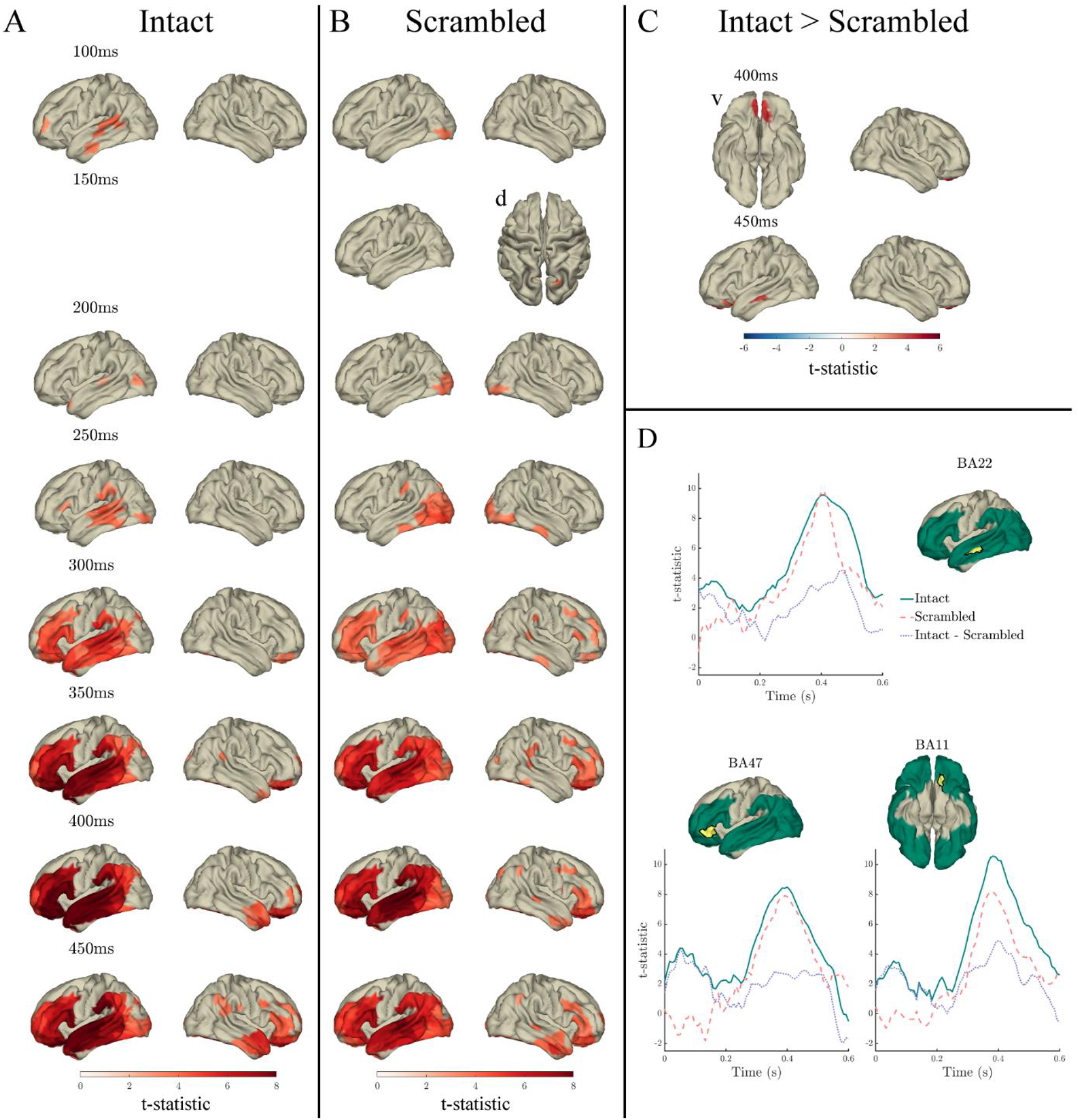
Effects of surprisal in the response to content words: Surface plots of T-statistics (averaged over 50ms time windows centred at the indicated latencies, for visualisation) quantifying the difference in variance explained by surprisal, beyond that explained by lexical frequency (log10 transformed), index, entropy and length, in intact sentence compared to random permutation models (panel A; *p*<.05 one-sided, corrected), scrambled sentence compared to random permutation models (panel B; *p*<.05 one-sided, corrected), and intact compared to scrambled sentence models (panel C; *p*<.05 two-sided, corrected). Parcels for which no time point was significant during the 50ms time bin are masked. Ventral and dorsal views are indicated with adjacent “v” and “d” labels, respectively. Panel D: Time courses of T-statistics for intact (solid green line) and scrambled (dashed red line) sentence models compared to random permutation models, and intact compared to scrambled sentence models (dotted purple line) for subparcels of BA22, BA47 and BA11 (highlighted in yellow on adjacent surface plots). ROIs entered into statistical analyses are illustrated as green shaded area on surface plots.

**Figure 6.**
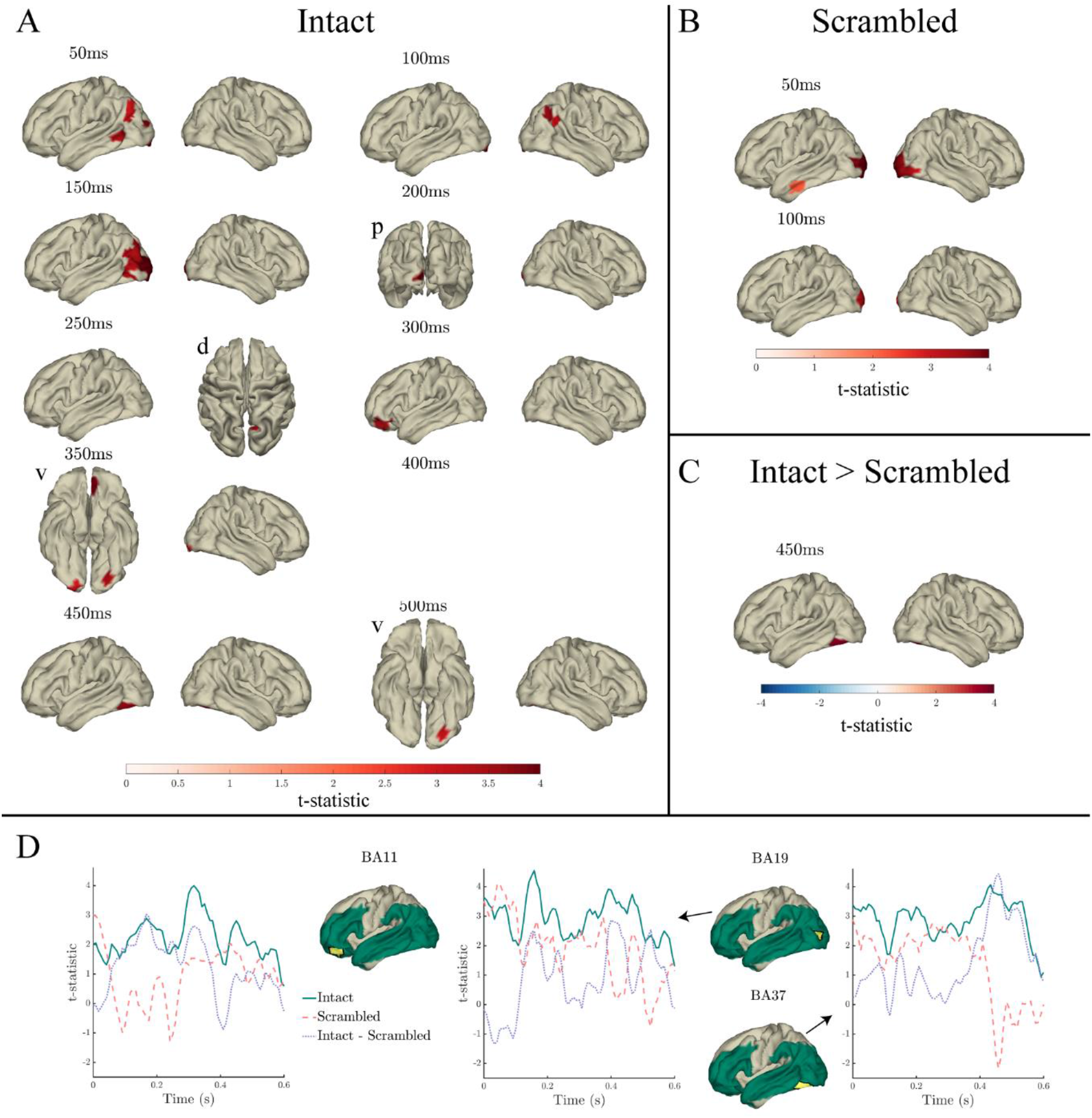
Effects of entropy in the response to content words: Surface plots of T-statistics (averaged over 50ms time windows centred at the indicated latencies, for visualisation) quantifying the difference in variance explained by entropy, beyond that explained by lexical frequency (log10 transformed), index, surprisal and length, in intact sentence compared to random permutation models (panel A; *p*<.05 one-sided, corrected), scrambled sentence compared to random permutation models (panel B; *p*<.05 one-sided, corrected), and intact compared to scrambled sentence models (panel C; *p*<.05 two-sided, corrected). Parcels for which no time point was significant during the 50ms time bin are masked. Ventral, dorsal, and posterior views are indicated with adjacent “v”, “d” and “p” labels, respectively. Panel D: Time courses of T-statistics for intact (solid green line) and scrambled (dashed red line) sentence models compared to random permutation models, and intact compared to scrambled sentence models (dotted purple line) for subparcels of BA11, BA19 and BA37 (highlighted in yellow on adjacent surface plots). ROIs entered into statistical analyses are illustrated as green shaded area on surface plots.

**Figure 7.**
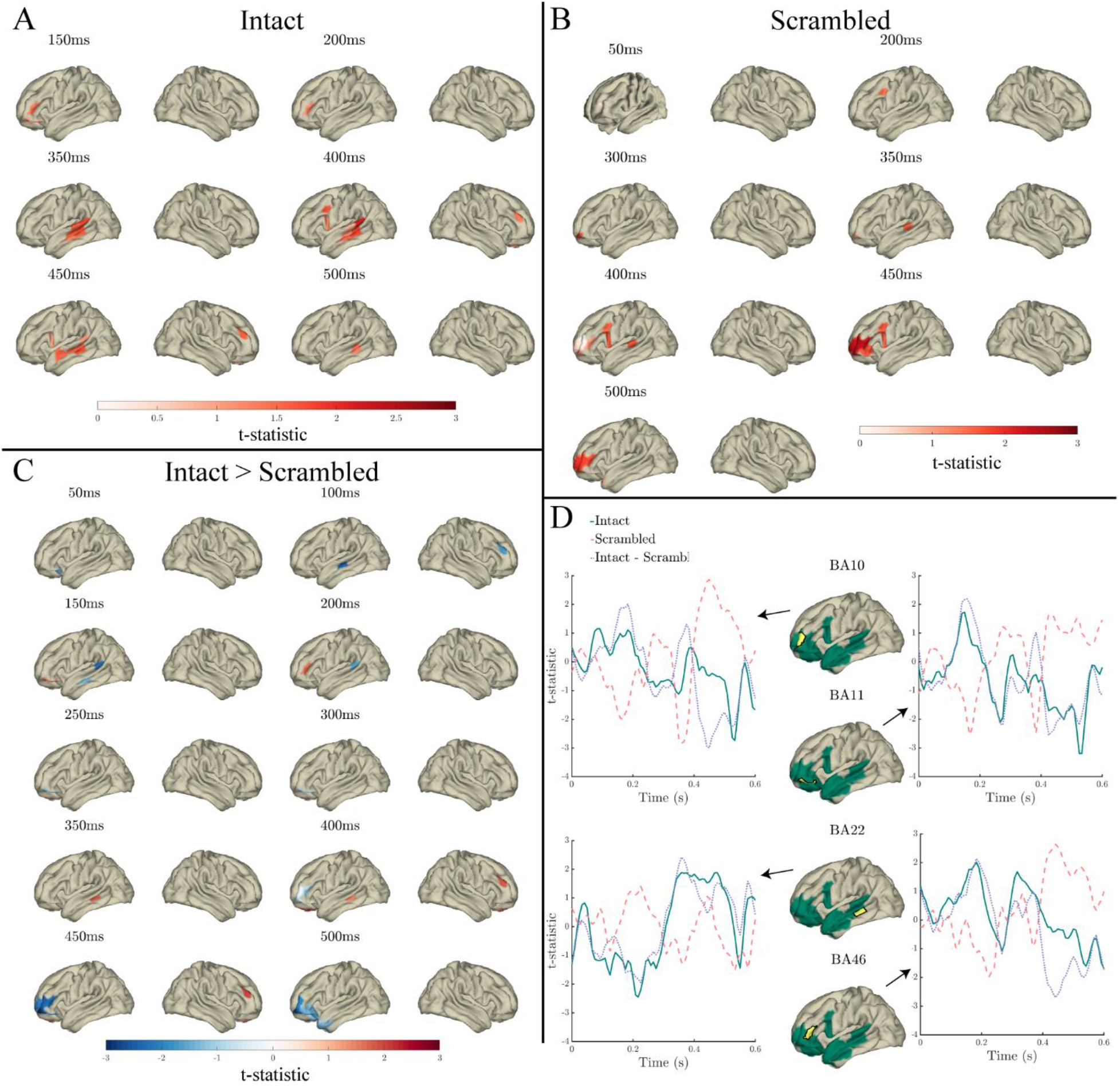
Effects of the lexical frequency × index interaction in the response to content words: Surface plots of T-statistics (averaged over 50ms time windows centred at the indicated latencies, for visualisation) quantifying the difference in variance explained by lexical frequency × index interaction, beyond that explained by lexical frequency (log10 transformed), index, and length in intact sentence compared to random permutation models (panel A; *p*<.05 one-sided, uncorrected), scrambled sentence compared to random permutation models (panel B; *p*<.05 one-sided, uncorrected), and intact compared to scrambled sentence models (panel C; *p*<.05 two-sided, corrected). Parcels for which no time point was significant during the 50ms time bin are masked. Panel D: Time courses of T-statistics for intact (solid green line) and scrambled (dashed red line) sentence models compared to random permutation models, and intact compared to scrambled sentence models (dotted purple line) for subparcels of BA44, BA22, BA10 and BA11 (highlighted in yellow on adjacent surface plots). ROIs entered into statistical analyses are illustrated as green shaded areas on surface plots.

**Figure 8.**
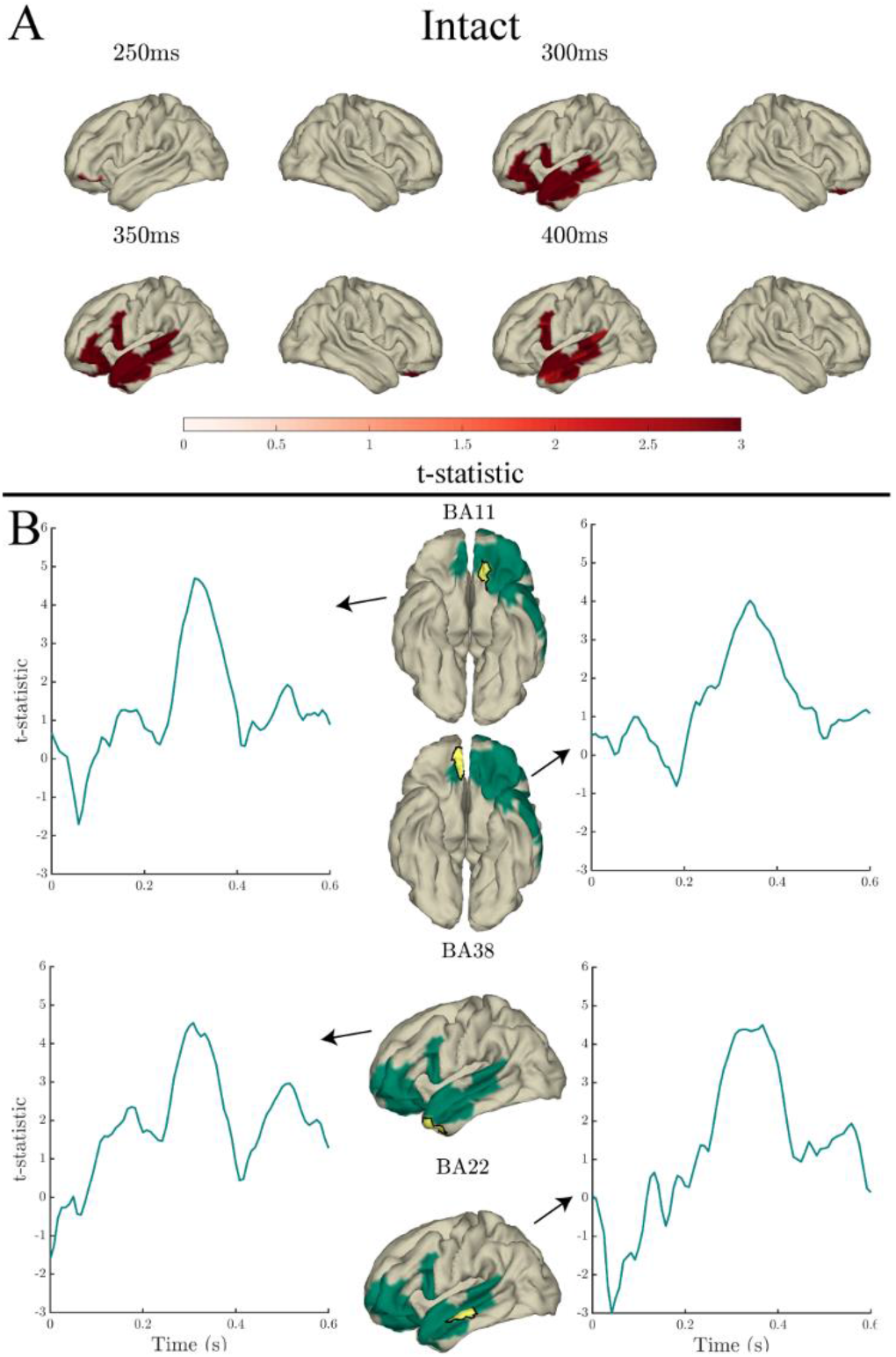
Effects of the lexical frequency × surprisal interaction in the response to content words: Surface plots of T-statistics (averaged over 50ms time windows centred at the indicated latencies, for visualisation) quantifying the difference in variance explained by lexical frequency × surprisal interaction, beyond that explained by lexical frequency (log10 transformed), surprisal (log10 transformed), and word length, in intact sentence compared to random permutation models (panel A; *p*<.05 one-sided, corrected). Parcels for which no time point was significant during the 50ms time bin are masked. Panel B: Time courses of T-statistics for intact sentences models compared to random permutation models, for subparcels of BA11 (left hemisphere), BA11 (right hemisphere), BA38 and BA22 (highlighted in yellow on adjacent surface plots). ROIs entered into statistical analyses are illustrated as green shaded areas on surface plots.

**Figure 9.**
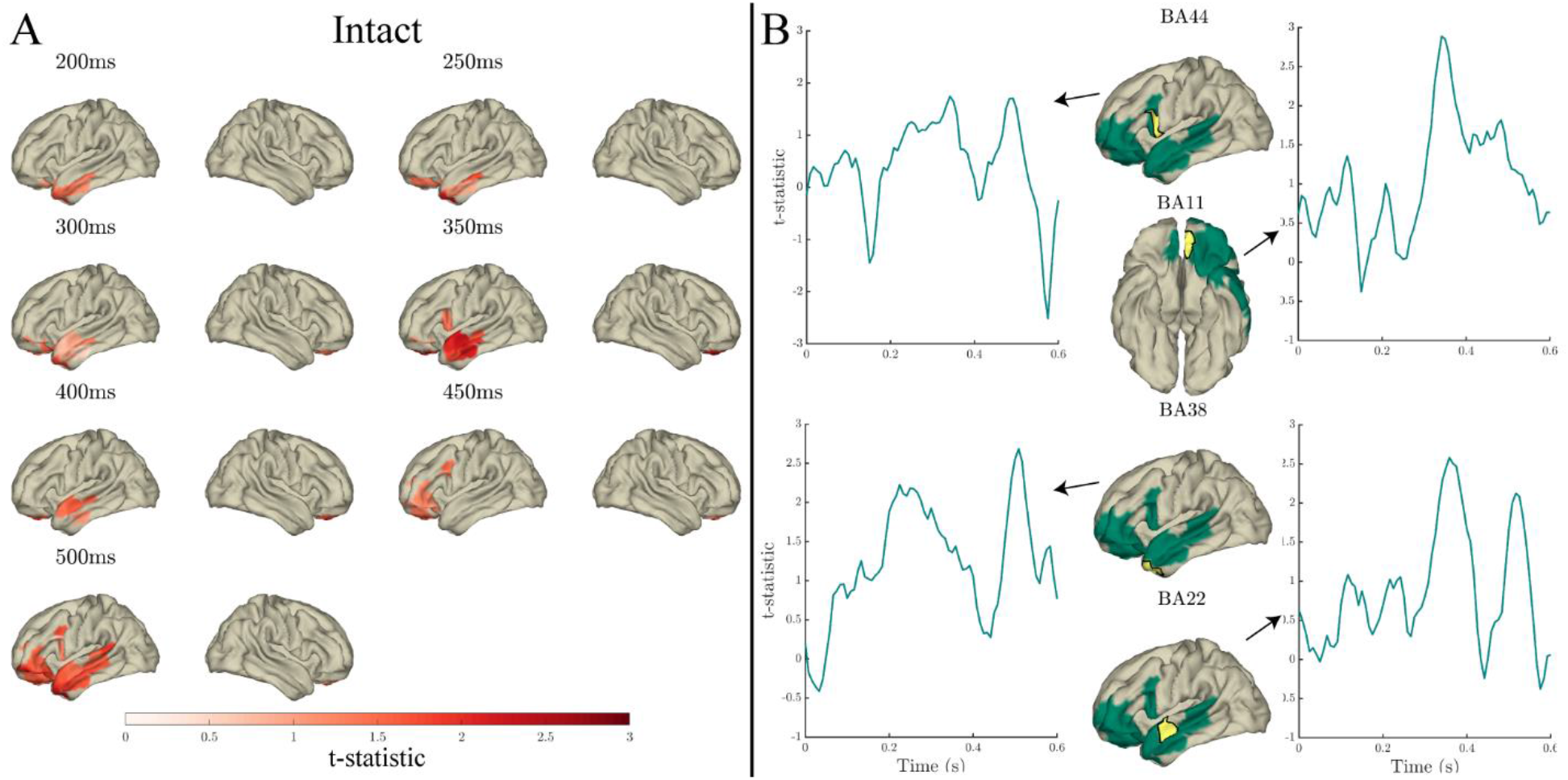
Effects of the lexical frequency × entropy interaction in the response to content words: Surface plots of T-statistics (averaged over 50ms time windows centred at the indicated latencies, for visualisation) quantifying the difference in variance explained by lexical frequency × entropy interaction, beyond that explained by lexical frequency (log10 transformed), entropy, and word length, in intact sentence compared to random permutation models (panel A; *p*<.05 one-sided, uncorrected). Parcels for which no time point was significant during the 50ms time bin are masked. Panel B: Time courses of T-statistics for intact sentences models compared to random permutation models, for subparcels of BA44, BA11, BA38 and BA22 (highlighted in yellow on adjacent surface plots). ROIs entered into statistical analyses are illustrated as green shaded areas on surface plots.

#### Lexical Frequency

Lexical frequency significantly predicted MEG signal in both intact and scrambled sentences throughout the 0-600ms analysis window (relative to word onset), spatially spreading from bilateral occipital and inferior temporal cortex to left posterior and middle temporal cortex at time points preceding 250ms, to left frontal and left anterior temporal cortex from 250ms onwards. In both intact and scrambled sentences, the effect of lexical frequency peaked at around 400ms in left temporal and frontal cortex (Fig 3 panels A and B). In the left superior temporal gyrus (STG) and middle temporal gyrus (MTG) this effect started earlier in intact compared to scrambled sentences, from 183ms, compared to 267ms in scrambled sentences.

Despite the seemingly stronger effect in scrambled compared to intact sentences -apparent in the time courses in Fig 3 panel D - in a direct comparison of the coefficient of determination for lexical frequency across conditions (presented in Fig 3 panel C), only a very small spatiotemporal effect survived the multiple comparisons correction scheme. Specifically, significantly more variance was explained in intact compared to scrambled sentences at a single time point, at 267ms, in a single right hemisphere frontal parcel (BA46). There were no other significant differences between intact and scrambled sentences in the variance explained by lexical frequency (corrected *p*>.05).

#### Index

Index significantly predicted the MEG signal in both intact and scrambled sentences throughout the 0-600ms analysis window. In intact sentences the effect spread from bilateral occipital cortex throughout right posterior and inferior temporal cortex and left temporal and frontal cortex, and peaked at around 350ms in left anterior temporal and inferior frontal cortex (Fig 4 panel A). In contrast to intact sentences, in scrambled sentences the effect was predominantly constrained to bilateral occipital and inferior temporal cortex, peaked at around 300ms in left posterior and inferior temporal cortex (Fig 4 panel B), and after 492ms only two single time points were significant (542ms and 600ms).

Significantly more variance in the MEG signal was predicted by index in intact compared to scrambled sentences from 275-417ms in anterior temporal (BA21/22/38), 283-375ms in inferior frontal (BA44/46/47), and 258-400ms in orbitofrontal and prefrontal cortex (PFC; BA10/11), as is evident in Fig 4 panel C.

#### Surprisal

In both intact and scrambled sentences, surprisal significantly predicted the MEG signal throughout most of the analysis window, peaking at 400ms in temporal and frontal cortex, and predicting additional right hemisphere variance in orbitofrontal and anterior temporal cortex (see Fig 5 panels A and B). In intact sentences, the effect spread from STG and the angular gyrus from 0-100ms, throughout temporal and frontal cortex from 208-600ms (Fig 5 panel A). In scrambled sentences, the effect spread from left (later bilateral) occipital and inferior temporal cortex throughout primarily the left temporal and frontal cortex (Fig 5 panel B). However, the effect of surprisal in scrambled sentences was most robust from 200ms onwards. Preceding 200ms, only several individual time points were significant after multiple comparisons correction.

Significantly more variance in the MEG signal was predicted by surprisal in intact compared to scrambled sentences 50-58ms and 458-475ms relative to word onset in left MTG (BA22), and 392-442ms relative to word onset in bilateral orbitofrontal cortex (BA11), which is presented in Fig 5 panel C. The time courses in Fig 5 panel D illustrate that the significant difference in BA22 results from a more sustained response in intact compared to scrambled sentences, whereas BA11 results from a greater peak in intact compared to scrambled sentences.

#### Entropy

Entropy significantly predicted the MEG signal in both intact and scrambled sentences, however to a lesser extent than the aforementioned predictors. In intact sentences (see Fig 6 panel A), entropy predicted variance in bilateral occipital, left inferior temporal and frontal parcels, and in the posterior MTG, from 0-242ms, 292-367ms, and at individual time points of 433ms and 525ms (relative to word onset). In scrambled sentences (see Fig 6 panel B), entropy significantly predicted variance in bilateral occipital and inferior temporal cortex from 0-83ms. Significantly more variance was explained by entropy in intact compared to scrambled sentences in a single left inferior temporal parcel (BA37) 450-458ms after word onset (see Fig 6 panel C).

#### Condition specific interactions between lexical frequency and predictability

The above findings show how our model comparison approach identified brain activity patterns that were aligned with word-by-word fluctuations of various quantities that relate to lexical predictability. Considering that the interaction between lexical frequency and context (often quantified with word position in the sentence) has been consistently reported in previous electrophysiological studies (Alday et al., 2017; Dambacher et al., 2006; Payne et al., 2015; Sereno et al., 2019; Van Petten & Kutas, 1990), we conducted an analysis of this interaction in our data – in parcels that showed conditional differences in effects of either lexical frequency or index (see Figs 3-4) – specifically focussing on the spatial and temporal dynamics of this effect.

The interaction between lexical frequency and increased word position in the sentence is thought to occur through the increasingly constraining context facilitating predictability as the sentence progresses (Dambacher et al., 2006; Payne et al., 2015; Van Petten & Kutas, 1990). Indeed, as outlined in the Introduction, effects of lexical frequency and word predictability have been found to interact (Dambacher et al., 2012; Dambacher et al., 2006; Kretzschmar et al., 2015; Sereno et al., 2003; Sereno et al., 2019). Hence, in addition to investigating the interaction between lexical frequency and index, we conducted an analysis of the interaction between lexical frequency and measures of local predictability, surprisal and entropy. Given that there were only sparse differences between intact and scrambled sentences in the effects of surprisal and entropy, suggesting that surprisal and entropy quantify similar processing mechanisms regardless of the level of sentential context, these interactions were investigated in sentences only, and in the same parcels in which the lexical frequency × index interaction was investigated, in order to remain consistent across analyses.

Only effects of the interaction between lexical frequency and surprisal survived the stringent multiple comparisons correction (*p*<.05 corrected), and not the interactions with index and entropy. Fig 8 presents the spatiotemporal distributions, along with example time courses, of T-statistics for parcels and time points that were significant while correcting for multiple comparisons (*p*<.05 corrected), whereas Figs 7 and 9 present those that were significant without correcting for multiple comparisons (uncorrected *p*<.05).

##### Lexical frequency × Index

Beyond the variance explained by the main effects of index, lexical frequency, and length the index × lexical frequency interaction explained additional variance in intact sentences from 150ms after word onset in frontal parcels (BA10/BA11/BA44/BA47), spreading to MTG and posterior STG (BA22/BA38) from 342ms onwards, where effects peaked at around 400ms (see Fig 7 panel A; uncorrected *p*<.05).

In scrambled sentences, the index × lexical frequency interaction explained additional variance in several time windows throughout the 0-600ms analysis window, predominantly from 300ms onwards, but also at earlier time points. The effect spread from frontal (BA10/BA11) to temporal (BA22/BA38) and inferior frontal (BA44/BA46) parcels, peaking at around 450ms in frontal parcels (see Fig 7 panel B; uncorrected *p*<.05).

On inspection of Fig 7 panel C, the comparison of the coefficient of determination for the interaction in intact and scrambled sentence models revealed an interesting spatiotemporal pattern of results. During an earlier time window (100-300ms), more variance was explained by the index × lexical frequency interaction in intact compared to scrambled sentences in frontal parcels (BA10/BA11; warm colours Fig 7 panel C), yet more variance was explained by the index × lexical frequency interaction in scrambled compared to intact sentences in temporal parcels (BA21/BA22/BA38; cool colours Fig 7 panel C). However, in a later time window (350-500ms) a reverse pattern was observed, where more variance was explained by the interaction in scrambled compared to intact sentences in frontal parcels, and more variance was explained in intact compared to scrambled sentences in temporal parcels. This pattern is also evident in the time courses of T-statistics presented in Fig 7 panel D.

##### Lexical frequency × Surprisal

The interaction between lexical frequency and surprisal significantly predicted MEG signal variance, beyond the main effects of lexical frequency, surprisal and word length, from 275-392ms, starting in a frontal parcel (BA11), and spreading to anterior temporal parcels from 283ms (BA38), and further throughout temporal (BA21/BA22) and frontal (BA44/BA46) parcels, from 292ms and 308ms respectively. Effects peaked at around 350ms (see Fig 8 panel A-B; corrected *p*<.05).

##### Lexical frequency × Entropy

Beyond the variance explained by the main effects of entropy, lexical frequency, and length, the lexical frequency × entropy interaction explained additional variance from 200-600ms, starting in anterior temporal (BA38/BA21/BA22) and frontal (BA11) parcels, spreading further throughout frontal parcels (BA10/BA44) from 250ms/330ms (respectively) and posteriorly through middle and superior temporal cortex (see Fig 9; uncorrected *p*<.05). The effect of the interaction peaked at around 350ms, and again at approximately 500ms.

## 4. Discussion

During sentence reading, the brain processes individual words at a remarkable speed. Such fast processing is not only facilitated and affected by the word’s frequency of occurrence within a given language (Calvo & Meseguer, 2002; Inhoff & Rayner, 1986; Rayner & Duffy, 1986; Rubenstein et al., 1970), but also by the word’s context, brought about by semantic and syntactic constraints imposed by preceding words (Calvo & Meseguer, 2002; Staub et al., 2015; Van Petten & Kutas, 1990). There is a well-documented discrepancy between the electrophysiological and eye-tracking literature as to whether frequency and context have additive or interactive effects on processing (Kretzschmar et al., 2015). It is unclear whether word frequency influences processing when the input is predictable. The current work aimed to better define the spatiotemporal dynamics of the effects of lexical frequency and predictability on word processing, establish to what extent lexical frequency and predictability independently influence word processing, and to what extent they interact. To this end, we performed state-of-the-art analysis of a large and well-balanced MEG dataset, combining spatiotemporal hyperalignment with cross-validated encoding model comparisons. This allowed us to go beyond the more traditional approaches that use event-related averaging or generalized linear models, thus being able to infer effects based on the brain’s response to individual words.

We found that the MEG signal reflects the lexical frequency of individual words (here content words) throughout the analysis time window beyond effects of predictability, in a network expanding from occipital cortex throughout the left temporal and inferior frontal regions of the language network. Index, surprisal, and entropy additionally each significantly predicted the MEG signal. All comparisons were made while controlling for each alternative predictor, and word length. There were significant but focal differences between intact and scrambled sentences in the effects of lexical frequency, surprisal and entropy. In contrast to these focal differences, the effect of index differed extensively in intact compared to scrambled sentences. Thus, out of the analysed predictors, only the effect of index was greatly influenced by the sentential context in which words were presented (i.e. intact/scrambled sentences). These findings highlight that the word processing mechanisms reflected by index are dependent on the preceding context, whereas the processing mechanisms underlying lexical frequency and surprisal remain largely the same regardless of the degree of sentential context. Finally, only the interaction between lexical frequency and surprisal survived multiple comparisons correction (in ventromedial PFC and anterior temporal lobe), and not the interaction between lexical frequency and entropy, nor between lexical frequency and index. Although the index × lexical frequency interaction effect was not significant under a conservative multiple comparisons correction scheme, an inspection of the uncorrected results uncovered an interesting pattern. Namely, both left temporal and frontal cortical activity seemed to be influenced by the interaction, yet the latency at which this occurred was flipped across conditions. While, in intact sentences, the interaction was expressed more strongly at early time points in frontal areas and only later in temporal areas, this pattern was reversed for scrambled sentences. Importantly, on inspection of both the corrected and uncorrected results, the interactions between lexical frequency and our metrics quantifying predictability show an initial peak between 150-250ms. Given that the average fixation duration lasts ∼200ms, any processing related to eye movement decisions must occur prior to this time window (Sereno & Rayner, 2003). Our findings tentatively support that lexical frequency and predictability do not interact robustly until around 150ms or later, which could explain why eye movement studies display a purely additive effect of these variables, in contrast to the robust interaction observed across electrophysiological studies. In the following paragraphs we discuss the results in more detail.

### 4.1. Lexical frequency

Overall, lexical frequency was encoded in the MEG signal to a similar extent in intact and scrambled sentences. This effect was widespread, both in space and time, and thus suggests that lexical frequency generically affects the brain response, likely reflecting less effortful processing of high compared to low frequency words. These findings help to close the gap between the electrophysiological and eye tracking literature, by providing evidence that frequency indeed influences word processing independently of prediction. In contrast to the eye tracking literature, electrophysiological studies have previously found that, during word processing, effects of lexical frequency disappear with increased context (Dambacher et al., 2006; Payne et al., 2015; Sereno et al., 2019; Van Petten & Kutas, 1990).

Although our findings differ from Fruchter et al. (2015), who found that word frequency explained no additional variance in the MEG signal after word onset beyond that explained by predictability, our results are consistent with the overall findings from the paper. Specifically, the authors presented evidence that, rather than reflecting a baseline level of predictability, lexical frequency influenced lexical access itself, as the frequency of the predicted word affected the electrophysiological response in the MTG prior to seeing the word (i.e. in response to the highly constraining word).

In the current data, effects of lexical frequency were observed after controlling for predictability prior to 100ms in occipital cortex. Such an early response in visual processing regions likely reflects an influence of word frequency on identification of the word form. To measure the extent that these early effects could be explained by the frequency of lower level sublexical properties of the word form, rather than the frequency of the lexeme, we conducted an additional analysis of lexical frequency while controlling for bigram and trigram letter frequency, as well as all other predictors (see Methods section 2.7). The results of this analysis are presented in the supplementary material (Fig SM1). Here it can be seen that the overall effect of word frequency remained the same as compared to when these variables were not controlled for (see Fig 3). Although lexical frequency explained variance in a reduced number of occipital and occipito-temporal parcels while controlling for the words’ lower level visual characteristics, compared to the results presented in Fig 3, an effect of lexical frequency was still observed in visual cortex at around 100ms. Lexical frequency therefore seems to influence early visual processing, beyond effects of the frequency of lower level properties of the word form.

The effect of lexical frequency progressively moved anteriorly through temporal and frontal cortex throughout word processing, supporting that lexical frequency influences multiple stages of word processing, such as lexical access and integration with the sentential context. These findings are in line with the EZ model of word reading (Reichle, Pollatsek, & Rayner, 2012), which proposes that word frequency and predictability independently affect both early (word form recognition) and late (lexical access/integration/compositional) stages of processing. Comparing intact and scrambled sentences, frequency was encoded in the MEG signal earlier in intact than scrambled sentences in the STG and MTG. Given the association of the MTG with lexical–semantic processing (Friederici, 2012; Hagoort, 2017) and the location of the primary auditory cortex and auditory association areas on the STG, the current results suggest that lexical frequency facilitates aspects of semantic and phonological processing earlier when the word is presented in a coherent sentence than when presented in a scrambled sentence. Moreover, significantly more variance was explained in intact compared to scrambled sentences at 267ms in a single dorsolateral PFC parcel (BA46), an area thought to be involved in executive control during language processing (Hagoort, 2003, 2013, 2017).

### 4.2. Sentential context and predictability

In line with previous literature (Armeni, Willems, van den Bosch, & Schoffelen, 2019; Hultén, Schoffelen, Uddén, Lam, & Hagoort, 2019; Schuster et al., 2020), the word-by-word association between the MEG signals and the increasingly constrained context (i.e. index), and metrics quantifying (the results of) prediction, presented itself with different spatiotemporal dynamics. These will be discussed in the following paragraphs.

#### Index

Index explained a significant portion of variance in the MEG signal during the entire critical window in both intact and scrambled sentences. Moreover, index predicted the MEG signal significantly more in intact than scrambled sentences, predominantly in anterior temporal and frontal cortex. This latter finding illustrates that it is the progressing sentential context that affects word processing in these regions, rather than more domain-general properties that correlate with index, such as working memory demands. The anterior temporal lobe has been associated with conceptual representations (Peelen & Caramazza, 2012; Pylkkänen, 2019; Ralph, Jefferies, Patterson, & Rogers, 2017; Rice, Lambon Ralph, & Hoffman, 2015) and syntactic structure building (Brennan et al., 2012; Brennan & Pylkkänen, 2017), the latter of which is engaged more when words are presented in intact compared to scrambled sentences. The greater influence of index in intact compared to scrambled sentences in the inferior frontal gyrus is consistent with the notion of unification, the integration of lexical items within the wider semantic and syntactic context as the sentence unfolds (Hagoort, 2005, 2013).

In line with earlier work (Schuster et al., 2020), index was encoded in the MTG and angular gyrus in intact sentences. No such effect was observed in these regions for scrambled sentences, although the latter qualitative difference was not significant when directly contrasting conditions. Given the association between MTG activity and lexical-semantic processing (Friederici, 2012; Hagoort, 2017), the effect in MTG could reflect the build-up of richer semantic representations as coherent sentences progress, more so than during the progression of scrambled sentences. The absence of an effect of index in scrambled sentences in the angular gyrus may be consistent with the view that this region is a hub to integrate different types of information extracted by various parts of the language network (Binder & Desai, 2011; Hagoort, 2003, 2019). Although, the precise roles of the angular gyrus and the anterior temporal lobe in integrating conceptual information are still currently debated (Binder & Desai, 2011; Hagoort, 2019; Matchin, Liao, Gaston, & Lau, 2019; Pylkkänen, 2019; Ralph et al., 2017). In contrast to unfolding well-formed sentences, scrambled sentences lack syntactic structure, and therefore do not permit for a meaningful integration of structural cues with, for instance, lexico-semantic information.

#### Surprisal

We estimated surprisal and entropy using corpus-based statistics, using a tri-gram model on the individual intact and scrambled sentences. Consistent with our expectations, surprisal was overall larger in scrambled sentence words (see Fig 1). Yet, aside from subtle differences between intact and scrambled sentences, as discussed below, the overall spatiotemporal characteristics of MEG signal variance explained by surprisal, on top of the other predictors, was similar between conditions. One tentative explanation for this could be that the inclusion of the index predictor in the ‘baseline model’ already accounted for a large part of signal variance (albeit to different degrees across conditions), causing the additional information provided by surprisal values to be less distinctive across conditions. The word-by-word fluctuations in surprisal explained widespread, predominantly left-lateralized, brain signals, irrespective of condition. This suggests a relation between our operationalisation of surprisal on the one hand, and more automatic ease-of-integration related processes on the other hand. Although care was taken to scramble sentences in a way so as no more than three consecutive words could be syntactically combined, there is evidence that combinatorial processes are robust to local word swaps (Mollica et al., 2020). In the current data, surprisal seems to reflect the same underlying combinatorial processes in scrambled and intact sentences, reflecting the ease-of-integration.

A direct statistical comparison across conditions showed some very focal and short-lived differences. Apart from a very early time window, at around 50ms in the MTG, there was a difference around 400-450ms in orbitofrontal and MTG parcels. It is often difficult to determine whether observed effects of surprisal result from participants predicting the upcoming linguistic input, or from more probable words being easier to integrate (Pickering & Gambi, 2018; Willems, Frank, Nijhof, Hagoort, & van den Bosch, 2016). While the early effect of surprisal that we observe here is likely related to predictive mechanisms, the later MEG signatures might equally be caused by hindered integration. Surprisal was encoded in the MEG signal in temporal cortex prior to 100ms (in the sentence condition only), which has previously been argued to imply that some linguistic information about a word has been pre-activated – here constrained by the previous two words – given that bottom-up lexical retrieval could not yet have taken place (Pickering & Gambi, 2018). Although the precise timing of lexical access of written words is currently debated, it is thought that sub-lexical characteristics and the word form have been processed by ∼100ms and morphemic processing and lexical access of the lemma occurs between 150-170ms (Grainger & Holcomb, 2009; Hauk et al., 2006; Lewis, Solomyak, & Marantz, 2011; Pulvermüller, Shtyrov, & Hauk, 2009; Sereno & Rayner, 2003; Woollams, 2015). Such timings speak to a pre-activation account of the early effects of surprisal in the temporal cortex here. Sentence context may influence the timing of lexical retrieval through prediction mechanisms (Fruchter et al., 2015). In contrast, the later effects of surprisal at 400-450ms in orbitofrontal and MTG parcels could result from either integrative or predictive processes. Although the orbitofrontal cortex (situated in the ventromedial PFC) has previously been sensitive to predictability of both linguistic information (Hofmann et al., 2014) and more generally (Nobre, Coull, Frith, & Mesulam, 1999), the ventromedial PFC has also been associated with higher level combinatorial processes (Brennan & Pylkkänen, 2008, 2010; Pylkkänen, 2008, 2019, 2020; Pylkkänen, Martin, McElree, & Smart, 2009; Pylkkänen & McElree, 2007), in line with an integrative account of the later effect of surprisal here.

#### Entropy

Entropy quantifies the uncertainty of the upcoming linguistic content (Pickering & Gambi, 2018; Willems et al., 2016). Entropy significantly predicted the MEG signal in both intact and scrambled sentences. Notably, the spatial and temporal extent of significant effects were much smaller than those of the other predictors. Here, entropy was encoded in early occipital cortical activity, both in intact and scrambled sentences. Additionally, in sentences, entropy effects were observed in left frontal cortex around 300ms, and in inferior temporal cortex around 450ms. Effects of prediction in occipital parcels during early time points have previously been used as evidence to support the notion that an active prediction of word form is employed by the brain (Dikker, Rabagliati, Farmer, & Pylkkänen, 2010; Pickering & Gambi, 2018). Rather than directly reflecting prediction, entropy quantifies the uncertainty about upcoming words (here based on the prior two words). Prediction of upcoming words was not possible in the scrambled sentence condition. Participants have been shown to quickly adapt their predictive behaviour to the predictability of the linguistic content of the current context (Bosker, van Os, Does, & van Bergen, 2019; Heyselaar, Peeters, & Hagoort, 2020; Thacker, Chambers, & Graham, 2018). It therefore seems unlikely that, when reading scrambled sentences, participants still pre-activated word forms that would usually be likely candidates to follow in a sentence. An alternative explanation for the early occipital cortical activity here is that, under uncertainty of upcoming linguistic input, more weight is placed on bottom-up (as opposed to top-down) signal, and more resources are allocated to visual processing. In contrast to the more generic interpretation of early entropy effects in visual cortical areas, the later sentence-specific effect in inferior temporal cortex could indeed reflect predictive processing of the word form. This region, often referred to as the visual word form area, is likely to receive top-down signals containing linguistic information about a word (Price & Devlin, 2011; Sharoh et al., 2019).

Entropy presented with a markedly different pattern of results compared to the other prediction metrics, in that only several focal groups of parcels during narrow time points survived multiple comparisons correction. It is evident from the time courses in Fig 6 that the encoding of the MEG signal was temporally less consistent for the entropy models compared to the models presented in Figs 3-5. Similarly, Schuster et al. (2020) found no effect of predictability (entropy) in the haemodynamic response when conducting a whole-brain analysis, and effects were found only in an ROI analysis.

### 4.3. Interactions between lexical frequency and predictability

In line with previous work (Alday et al., 2017; Dambacher et al., 2006; Fruchter et al., 2015; Payne et al., 2015; Sereno et al., 2019; Van Petten & Kutas, 1990), we investigated the interaction between lexical frequency and metrics quantifying prediction, including index (both within and across individual conditions), surprisal and entropy (in sentences only). Here we add to the previous literature by investigating the spatiotemporal dynamics of the interaction in more detail in comparison to previous reports (Fruchter et al., 2015). Using a strict multiple comparisons correction scheme, we found evidence of an interaction only between lexical frequency and surprisal, and not between lexical frequency and index, nor lexical frequency and entropy. The latter two findings seem to concur with the eye-tracking literature, which has found an additive effect of lexical frequency and predictability on fixation durations (Kennedy et al., 2013; Kretzschmar et al., 2015; Staub, 2015; Staub & Benatar, 2013). Yet, the lexical frequency × surprisal interaction results are in line with the electrophysiological literature, in which effects of lexical frequency on word processing are reduced with increased predictability (Dambacher et al., 2012; Dambacher et al., 2006; Kretzschmar et al., 2015; Sereno et al., 2003; Sereno et al., 2019). Furthermore, partially supporting the aforementioned electrophysiological literature, an analysis of the nominally thresholded data revealed a spatially similar pattern of results of the entropy interaction as compared to the significant interaction with surprisal (corrected *p*<.05), in addition to some interesting condition-specific dynamics of the lexical frequency × index interaction. Finally, all three interactions first peaked between 150-250ms, suggesting that these variables could *additively* influence early stages of word processing prior to 150ms, but interact during later, post-lexical stages of word processing. Such findings help to explain why, in contrast to the electrophysiological literature, only an additive effect of these variables has been observed in the eye tracking literature. Given that an average fixation duration lasts ∼200ms (Rayner, 1986), eye movement decisions should only be influenced by information obtained in early stages of word processing (Sereno & Rayner, 2003).

Firstly, lexical frequency interacted with surprisal and entropy in frontal (predominantly in BA11) and anterior temporal parcels, the interaction being strongest at around 350ms. Both the anterior temporal lobe and BA11 have been proposed to be involved in combinatorial processes during sentence comprehension, the former in semantic (Binder & Desai, 2011; Brennan & Pylkkänen, 2017; Hagoort, 2019; Matchin, Liao, et al., 2019; Pylkkänen, 2019; Ralph et al., 2017) or syntactic (Brennan et al., 2012; Brennan & Pylkkänen, 2017) integration, and the latter in higher level compositional processing and inferring implicit meanings (Brennan & Pylkkänen, 2008, 2010; Pylkkänen, 2008, 2019, 2020; Pylkkänen et al., 2009; Pylkkänen & McElree, 2007). The frequency of a word may therefore become less relevant to its integration within the higher level sentential meaning when the same word is highly predictable. Although we do not report the direction of the interaction here (see Section 4.4. Limitations and future work), previous reports have consistently shown that the effect of frequency on word processing diminishes with increased predictability, and the benefits of predictability on word processing are enhanced for low compared to high frequency words (Dambacher et al., 2012; Dambacher et al., 2006; Fruchter et al., 2015; Hofmann et al., 2014; Kretzschmar et al., 2015; Sereno et al., 2003; Sereno et al., 2019). For example, similar to the current results, an interaction between lexical frequency and predictability was found in orbitofrontal cortex (encompassed in BA11) by Hofmann et al. (2014), who found stronger brain responses to disconfirmed predictions for only low and not high frequency words.

The interaction between lexical frequency and index displayed some intriguing dynamics in time and space across conditions (despite not surviving multiple comparisons corrections). In left temporal parcels (BA21/BA22/BA38), including the MTG, the interaction explained more variance in scrambled than intact sentences at early time points, and in intact compared to scrambled sentences in a later time window. The later (350-500ms) temporal cortex effect is consistent with previous electrophysiological literature that has averaged over central-parietal sensors in an N400 time window, as the interaction explained more variance in coherent sentences than in scrambled sentences. Specifically, earlier work has shown that the effect of frequency on the N400 diminishes with increased word position, in intact sentences but not scrambled sentences (Payne et al., 2015), eliciting the conclusion that lexical frequency no longer influences word processing when there is increased context. An interaction between word frequency and predictability in the left MTG is also consistent with the findings of Fruchter et al. (2015), who found an effect of frequency here only for words of low and not high predictability. One mechanism through which this could occur is through the pre-activation of semantic features associated with the lexical item, or pre-activation of the lexical item itself, so that processing low frequency words is no longer as difficult compared to high frequency words.

In frontal parcels (BA10/BA11), more variance was explained by the interaction in intact compared to scrambled sentences in an early time window, and in scrambled compared to intact sentences in a later time window. Greater ventromedial PFC (BA11) recruitment has previously been observed in sentences compared to word-lists more generally (Brennan & Pylkkänen, 2012) and, as discussed above, is thought to be involved in interpreting higher level sentence meanings (Brennan & Pylkkänen, 2008, 2010; Pylkkänen, 2008, 2019, 2020; Pylkkänen et al., 2009; Pylkkänen & McElree, 2007). BA10, on the other hand, has been associated with encoding semantic relationships (Bunge et al., 2009; Frankland & Greene, 2020; Knowlton et al., 2012; Ramnani & Owen, 2004). Both higher level compositional processing and forming semantic relationships could be expected to occur earlier in intact compared to scrambled sentences. Overall, the difference between intact and scrambled sentences in the interaction between lexical frequency and index seems to occur in the time that these factors interact, rather than in the presence of an interaction.

Current models of word reading do not yet account for the effects observed in the current data, together with the aforementioned eye tracking and electrophysiological literature. The EZ-Reader model (Reichle et al., 2012), and more recent Uber-Reader model (Veldre, Yu, Andrews, & Reichle, 2020) of word reading, propose that lexical frequency and predictability independently influence both early (L1/identification of the word form) and late (L2/lexical access/semantic processing/integration) stages of word processing. While we provide confirmatory evidence that lexical frequency and predictability indeed influence both early and late stages of word processing independently, we also show that they interact during later stages of word processing. Models of word reading could therefore benefit from incorporating these additional findings. Although we do not quantify the direction of the interaction here, previous reports have robustly demonstrated that the effect of lexical frequency is reduced for highly predictable words (compare to unpredictable words), and the effect of predictability is greater for low than high frequency words (Dambacher et al., 2012; Dambacher et al., 2006; Fruchter et al., 2015; Hofmann et al., 2014; Kretzschmar et al., 2015; Sereno et al., 2003; Sereno et al., 2019).

### 4.4. Limitations and future work

A limitation of the current work is that words were presented word-by-word, causing the stimulation to be externally paced. Yet, it is well known that in more naturalistic settings the reading pace is determined by the reader, where eye movement and fixation behaviour is in part the result of prediction related processes (Rayner & Well, 1996). Indeed, there is evidence to suggest that predictability facilitates processing before a word is fixated, while the word is within parafoveal view (Balota, Pollatsek, & Rayner, 1985; Staub, 2015; Staub & Goddard, 2019). Furthermore, self-paced reading paradigms have demonstrated that fixation durations of the current word are influenced by the properties of the preceding word (Dambacher & Kliegl, 2007; Kliegl, Nuthmann, & Engbert, 2006). Predictive processes may be engaged at different latencies or to a different extent in natural reading compared to the current paradigm, due to their interaction with executive control of eye movements. Future work should aim to investigate whether the observed spatiotemporal dynamics of the effects of lexical frequency and predictability on the MEG signal hold during naturalistic reading (see Brennan & Pylkkänen, 2017 for an investigation into naturalistic word reading with MEG).

A further limitation of the current investigation is that we did not report the direction of the interaction between lexical frequency and our metrics quantifying predictability. As the variables in the models were highly correlated (see Fig 1), it is not possible to meaningfully interpret the beta weights from the models. Instead, we used a model comparison scheme to quantify the additional variance explained by each regressor, beyond that already explained by all other regressors (see Methods section 2.7 for details). Given that the direction of the interaction is robust across numerous previous reports, a lack of directionality in the current results does not greatly hinder the interpretation of our results. Moreover, by comparing intact and scrambled sentences, we were able to report the degree to which the strength of the interaction changed with and without sentential context. In doing so, we replicated previous findings of a stronger interaction in intact compared to scrambled sentences during a typical N400 time window in the temporal cortex. However, we additionally showed that the direction of this higher order interaction reversed in the frontal cortex during the same time window, and in the temporal cortex in an earlier time window. Future work with more carefully controlled stimuli could aim to replicate these results in a ROI analysis.

### 4.5. Conclusions

We provide evidence to support that frequency and contextual constraints have identifiable effects on multiple stages of word-by-word processing, from early visual and lexical retrieval to later integration and unification processes. Largely similar spatiotemporal effects across both intact and scrambled sentences suggest that lexical frequency generally affects how fast and effortful processing is, independently from ongoing predictive processes.

Only the interaction between lexical frequency and surprisal survived our conservative multiple comparisons corrections (in anterior temporal and frontal cortex), and not the interaction between lexical frequency and entropy, nor between lexical frequency and index. Although we found no significant effect of a lexical frequency × index interaction - consistent with the additive effects of these variables typically observed in the eye-tracking literature - an uncorrected analysis revealed some interesting spatiotemporal dynamics. Namely, the effect of the interaction was reversed in time and space in intact sentences compared to scrambled sentences. In the MTG, which is associated with lexical-semantic processing, the interaction explained more variance in scrambled sentences than intact sentences in an early time window, and in intact sentences than scrambled sentences in a later time window. The latter is consistent with the frequency × index interaction that is typically observed in the N400 time window in intact sentences but not scrambled sentences (Payne et al., 2015). In orbitofrontal and ventromedial PFC cortex, which have previously been associated with forming higher level semantic relationships and inferring implicit meanings, the interaction explained more variance in intact sentences than scrambled sentences at early time points, but in scrambled sentences than intact sentences in a later time window. Finally, we provide evidence to suggest that lexical frequency and predictability may independently influence early and late stages of word processing, but also interact during later stages of word processing. Our findings may contribute to improved models of word reading, which do not yet fully account for effects of predictability in the current results, nor in previous work (Staub & Goddard, 2019).

## Acknowledgements

We would like to thank Alessandro Lopopolo for computing the corpus-derived lexical characteristics of lexical frequency, surprisal and entropy for the current stimulus set.

## 6. Supplementary material

**Figure SM1.**
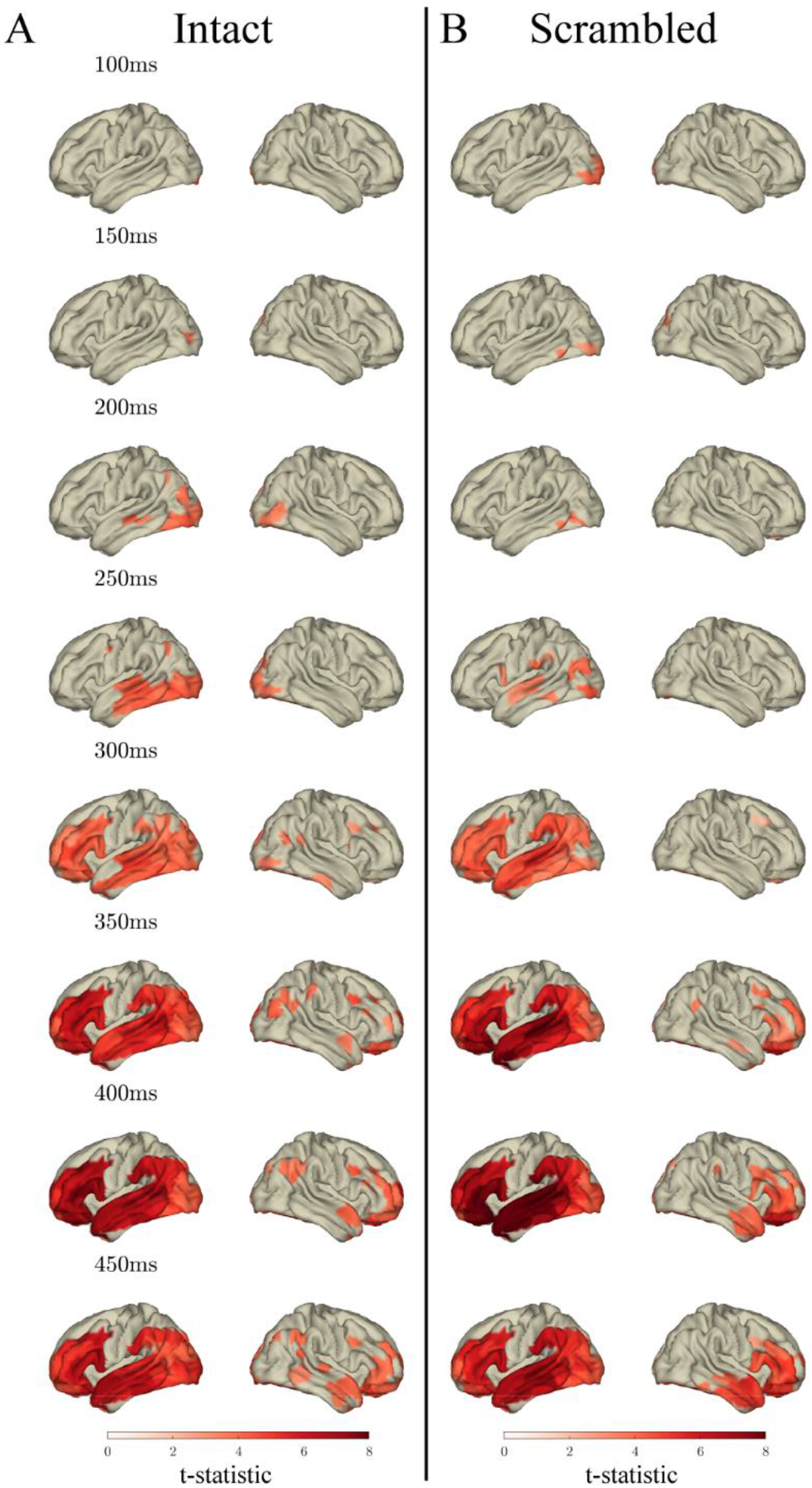
Effects of lexical frequency in the response to content words: Surface plots of T-statistics (averaged over 50ms time windows centred at the indicated latencies, for visualisation) quantifying the difference in variance explained by lexical frequency (log10 transformed), beyond that explained by index, surprisal (log10 transformed), entropy, length, bigram letter frequency (log10 transformed) and trigram letter frequency (log10 transformed) in intact sentence compared to random permutation models (panel A; *p*<.05 one-sided, corrected) and scrambled sentence compared to random permutation models (panel B; *p*<.05 one-sided, corrected). Parcels for which no time point was significant during the 50ms time bin are masked.

**Figure SM2.**
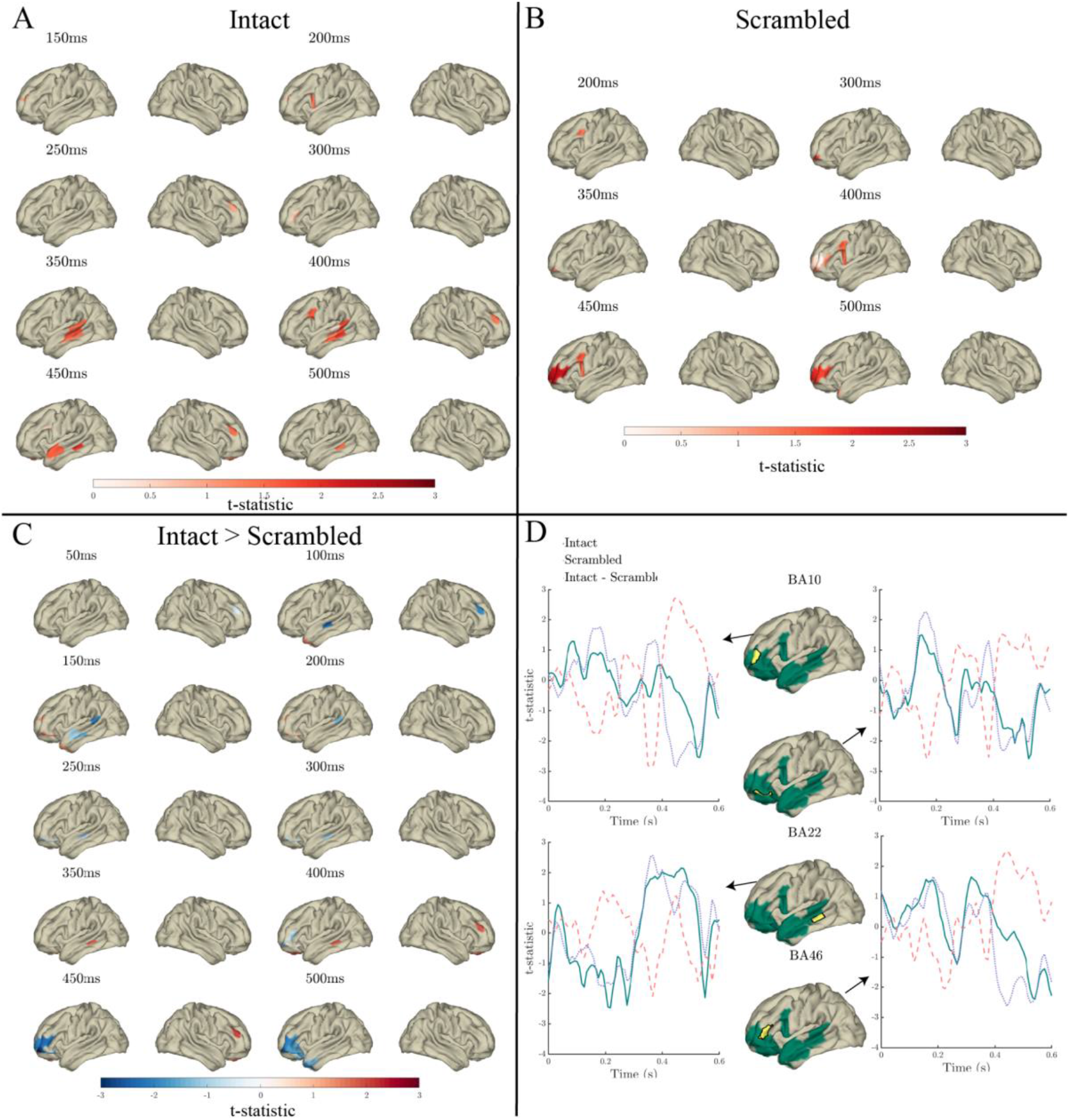
Effects of the lexical frequency × index interaction in the response to content words: Surface plots of T-statistics (averaged over 50ms time windows centred at the indicated latencies, for visualisation) quantifying the difference in variance explained by lexical frequency × index interaction, beyond that explained by lexical frequency (log10 transformed), index, length, bigram letter frequency (log10 transformed) and trigram letter frequency (log10 transformed) in intact sentence compared to random permutation models (panel A; *p*<.05 one-sided, uncorrected), scrambled sentence compared to random permutation models (panel B; *p*<.05 one-sided, uncorrected), and intact compared to scrambled sentence models (panel C; *p*<.05 two-sided, corrected). Parcels for which no time point was significant during the 50ms time bin are masked. Panel D: Time courses of T-statistics for intact (solid green line) and scrambled (dashed red line) sentence models compared to random permutation models, and intact compared to scrambled sentence models (dotted purple line) for subparcels of BA10, BA11 BA22 and BA46 (highlighted in yellow on adjacent surface plots). ROIs entered into statistical analyses are illustrated as green shaded areas on surface plots.

## References

Alday, P. M., Schlesewsky, M., & Bornkessel-Schlesewsky, I. (2017). Electrophysiology Reveals the Neural Dynamics of Naturalistic Auditory Language Processing: Event-Related Potentials Reflect Continuous Model Updates. eNeuro, 4(6). doi:10.1523/ENEURO.0311-16.2017

Arana, S., Marquand, A., Hultén, A., Hagoort, P., & Schoffelen, J.-M. (2020). Sensory Modality-Independent Activation of the Brain Network for Language. The Journal of Neuroscience, 40(14), 2914–2924. doi:10.1523/jneurosci.2271-19.2020

Armeni, K., Willems, R. M., van den Bosch, A., & Schoffelen, J.-M. (2019). Frequency-specific brain dynamics related to prediction during language comprehension. Neuroimage, 198, 283–295. doi:https://doi.org/10.1016/j.neuroimage.2019.04.083

Balota, D. A., Pollatsek, A., & Rayner, K. (1985). The interaction of contextual constraints and parafoveal visual information in reading. Cognitive Psychology, 17(3), 364–390. doi:https://doi.org/10.1016/0010-0285(85)90013-1

Binder, J. R., & Desai, R. H. (2011). The neurobiology of semantic memory. Trends in Cognitive Sciences, 15(11), 527–536. doi:https://doi.org/10.1016/j.tics.2011.10.001

Bosker, H. R., van Os, M., Does, R., & van Bergen, G. (2019). Counting ‘uhm’s: How tracking the distribution of native and non-native disfluencies influences online language comprehension. Journal of Memory and Language, 106, 189–202. doi:https://doi.org/10.1016/j.jml.2019.02.006

Brennan, J., Nir, Y., Hasson, U., Malach, R., Heeger, D. J., & Pylkkänen, L. (2012). Syntactic structure building in the anterior temporal lobe during natural story listening. Brain and Language, 120(2), 163–173. doi:https://doi.org/10.1016/j.bandl.2010.04.002

Brennan, J., & Pylkkänen, L. (2008). Processing events: Behavioral and neuromagnetic correlates of Aspectual Coercion. Brain and Language, 106(2), 132–143. doi:https://doi.org/10.1016/j.bandl.2008.04.003

Brennan, J., & Pylkkänen, L. (2010). Processing psych verbs: Behavioural and MEG measures of two different types of semantic complexity. Language and Cognitive Processes, 25(6), 777–807. doi:10.1080/01690961003616840

Brennan, J., & Pylkkänen, L. (2012). The time-course and spatial distribution of brain activity associated with sentence processing. Neuroimage, 60(2), 1139–1148. doi:https://doi.org/10.1016/j.neuroimage.2012.01.030

Brennan, J., & Pylkkänen, L. (2017). MEG Evidence for Incremental Sentence Composition in the Anterior Temporal Lobe. Cognitive Science, 41(S6), 1515–1531. doi:https://doi.org/10.1111/cogs.12445

Brysbaert, M. (2019). How many words do we read per minute? A review and meta-analysis of reading rate. Journal of Memory and Language, 109. doi:https://doi.org/10.1016h/j.jml.2019.104047

Bunge, S. A., Helskog, E. H., & Wendelken, C. (2009). Left, but not right, rostrolateral prefrontal cortex meets a stringent test of the relational integration hypothesis. Neuroimage, 46(1), 338–342. doi:https://doi.org/10.1016/j.neuroimage.2009.01.064

Calvo, M. G., & Meseguer, E. (2002). Eye Movements and Processing Stages in Reading: Relative Contribution of Visual, Lexical, and Contextual Factors. The Spanish Journal of Psychology, 5(1), 66–77. doi:10.1017/S1138741600005849

Catani, M., Allin, M. P. G., Husain, M., Pugliese, L., Mesulam, M. M., Murray, R. M., & Jones, D. K. (2007). Symmetries in human brain language pathways correlate with verbal recall. Proceedings of the National Academy of Sciences, 104(43), 17163–17168. doi:10.1073/pnas.0702116104

Chee, M. W. L., Hon, N. H. H., Caplan, D., Lee, H. L., & Goh, J. (2002). Frequency of Concrete Words Modulates Prefrontal Activation during Semantic Judgments. Neuroimage, 16(1), 259–268. doi:https://doi.org/10.1006/nimg.2002.1061

Chen, Y., Davis, M. H., Pulvermüller, F., & Hauk, O. (2015). Early Visual Word Processing Is Flexible: Evidence from Spatiotemporal Brain Dynamics. J Cogn Neurosci, 27(9), 1738–1751. doi:10.1162/jocn_a_00815

Crepaldi, D., Rastle, K., Coltheart, M., & Nickels, L. (2010). ‘Fell’primes ‘fall’, but does ‘bell’prime ‘ball’? Masked priming with irregularly-inflected primes. Journal of Memory and Language, 63(1), 83–99.

Dambacher, M., Dimigen, O., Braun, M., Wille, K., Jacobs, A. M., & Kliegl, R. (2012). Stimulus onset asynchrony and the timeline of word recognition: Event-related potentials during sentence reading. Neuropsychologia, 50(8), 1852–1870. doi:https://doi.org/10.1016/j.neuropsychologia.2012.04.011

Dambacher, M., & Kliegl, R. (2007). Synchronizing timelines: Relations between fixation durations and N400 amplitudes during sentence reading. Brain Research, 1155, 147–162. doi:https://doi.org/10.1016/j.brainres.2007.04.027

Dambacher, M., Kliegl, R., Hofmann, M., & Jacobs, A. M. (2006). Frequency and predictability effects on event-related potentials during reading. Brain Research, 1084(1), 89–103. doi:https://doi.org/10.1016/j.brainres.2006.02.010

de Cheveigné, A., Di Liberto, G. M., Arzounian, D., Wong, D. D. E., Hjortkjær, J., Fuglsang, S., & Parra, L. C. (2019). Multiway canonical correlation analysis of brain data. Neuroimage, 186, 728–740. doi:https://doi.org/10.1016/j.neuroimage.2018.11.026

Dikker, S., Rabagliati, H., Farmer, T. A., & Pylkkänen, L. (2010). Early Occipital Sensitivity to Syntactic Category Is Based on Form Typicality. Psychological Science, 21(5), 629–634. doi:10.1177/0956797610367751

Engbert, R., Nuthmann, A., Richter, E. M., & Kliegl, R. (2005). SWIFT: A Dynamical Model of Saccade Generation During Reading. Psychological Review, 112(4), 777–813. doi:10.1037/0033-295X.112.4.777

Frankland, S. M., & Greene, J. D. (2020). Two Ways to Build a Thought: Distinct Forms of Compositional Semantic Representation across Brain Regions. Cerebral Cortex, 30(6), 3838–3855. doi:10.1093/cercor/bhaa001

Friederici, A. D. (2009). Pathways to language: fiber tracts in the human brain. Trends in Cognitive Sciences, 13(4), 175–181. doi:https://doi.org/10.1016/j.tics.2009.01.001

Friederici, A. D. (2012). The cortical language circuit: from auditory perception to sentence comprehension. Trends in Cognitive Sciences, 16(5), 262–268. doi:https://doi.org/10.1016/j.tics.2012.04.001

Fruchter, J., Linzen, T., Westerlund, M., & Marantz, A. (2015). Lexical Preactivation in Basic Linguistic Phrases. Journal of Cognitive Neuroscience, 27(10), 1912–1935. doi:10.1162/jocn_a_00822

Glasser, M. F., & Rilling, J. K. (2008). DTI Tractography of the Human Brain’s Language Pathways. Cerebral Cortex, 18(11), 2471–2482. doi:10.1093/cercor/bhn011

Grainger, J., & Holcomb, P. J. (2009). Watching the Word Go by: On the Time-course of Component Processes in Visual Word Recognition. Language and Linguistics Compass, 3(1), 128–156. doi:10.1111/j.1749-818X.2008.00121.x

Hagoort, P. (2003). How the brain solves the binding problem for language: a neurocomputational model of syntactic processing. Neuroimage, 20, S18–S29. doi:https://doi.org/10.1016/j.neuroimage.2003.09.013

Hagoort, P. (2005). On Broca, brain, and binding: a new framework. Trends in Cognitive Sciences, 9(9), 416–423. doi:https://doi.org/10.1016/j.tics.2005.07.004

Hagoort, P. (2013). MUC (Memory, Unification, Control) and beyond. Frontiers in Psychology, 4(416). doi:10.3389/fpsyg.2013.00416

Hagoort, P. (2017). The core and beyond in the language-ready brain. Neuroscience & Biobehavioral Reviews.

Hagoort, P. (2019). The meaning-making mechanism(s) behind the eyes and between the ears. Philosophical Transactions of the Royal Society B: Biological Sciences, 375(1791). doi:doi:10.1098/rstb.2019.0301

Hagoort, P., Baggio, G., & Willems, R. M. (2009). Semantic unification. In The cognitive neurosciences, 4th ed. (pp. 819–836): MIT press.

Hauk, O., Patterson, K., Woollams, A., Watling, L., Pulvermüller, F., & Rogers, T. T. (2006). [Q:] When Would You Prefer a SOSSAGE to a SAUSAGE? [A:] At about 100 msec. ERP Correlates of Orthographic Typicality and Lexicality in Written Word Recognition. Journal of Cognitive Neuroscience, 18(5), 818–832. doi:10.1162/jocn.2006.18.5.818

Heyselaar, E., Peeters, D., & Hagoort, P. (2020). Do we predict upcoming speech content in naturalistic environments? Language, Cognition and Neuroscience, 1–22. doi:10.1080/23273798.2020.1859568

Hofmann, M. J., Dambacher, M., Jacobs, A. M., Kliegl, R., Radach, R., Kuchinke, L., … Herrmann, M. J. (2014). Occipital and orbitofrontal hemodynamics during naturally paced reading: An fNIRS study. Neuroimage, 94, 193–202. doi:https://doi.org/10.1016/j.neuroimage.2014.03.014

Huettig, F. (2015). Four central questions about prediction in language processing. Brain Research, 1626, 118–135. doi:https://doi.org/10.1016/j.brainres.2015.02.014

Hultén, A., Schoffelen, J.-M., Uddén, J., Lam, N. H. L., & Hagoort, P. (2019). How the brain makes sense beyond the processing of single words – An MEG study. Neuroimage, 186, 586–594. doi:https://doi.org/10.1016/j.neuroimage.2018.11.035

Inhoff, A. W., & Rayner, K. (1986). Parafoveal word processing during eye fixations in reading: Effects of word frequency. Perception & Psychophysics, 40(6), 431–439.

Kennedy, A., Pynte, J., Murray, W. S., & Paul, S.-A. (2013). Frequency and predictability effects in the Dundee Corpus: An eye movement analysis. The Quarterly Journal of Experimental Psychology, 66(3), 601–618. doi:10.1080/17470218.2012.676054

Kliegl, R., Nuthmann, A., & Engbert, R. (2006). Tracking the mind during reading: The influence of past, present, and future words on fixation durations. Journal of experimental psychology: general, 135(1), 12–35. doi:10.1037/0096-3445.135.1.12

Knowlton, B. J., Morrison, R. G., Hummel, J. E., & Holyoak, K. J. (2012). A neurocomputational system for relational reasoning. Trends in Cognitive Sciences, 16(7), 373–381. doi:https://doi.org/10.1016/j.tics.2012.06.002

Kresse, L., Kirschner, S., Dipper, S., & Belke, E. (2012). Towards exploring the specific influences of wordform frequency, lemma frequency and OLD20 on visual word recognition and reading aloud. Lexical Resources in Psycholinguistic Research, 3, 9.

Kretzschmar, F., Schlesewsky, M., & Staub, A. (2015). Dissociating Word Frequency and Predictability Effects in Reading: Evidence From Coregistration of Eye Movements and EEG. Journal of Experimental Psychology-Learning Memory and Cognition, 41(6), 1648–1662. doi:10.1037/xlm0000128

Kutas, M., & Federmeier, K. D. (2011). Thirty Years and Counting: Finding Meaning in the N400 Component of the Event-Related Brain Potential (ERP). Annual Review of Psychology, 62(1), 621–647. doi:10.1146/annurev.psych.093008.131123

Lam, N. H. L., Schoffelen, J. M., Udden, J., Hulten, A., & Hagoort, P. (2016). Neural activity during sentence processing as reflected in theta, alpha, beta, and gamma oscillations. Neuroimage, 142, 43–54. doi:10.1016/j.neuroimage.2016.03.007

Lau, E. F., & Namyst, A. (2019). fMRI evidence that left posterior temporal cortex contributes to N400 effects of predictability independent of congruity. Brain and Language, 199, 104697. doi:https://doi.org/10.1016/j.bandl.2019.104697

Lau, E. F., Phillips, C., & Poeppel, D. (2008). A cortical network for semantics: (de)constructing the N400. Nature Reviews Neuroscience, 9(12), 920–933. doi:10.1038/nrn2532

Levy, R. (2008). Expectation-based syntactic comprehension. Cognition, 106(3), 1126–1177. doi:https://doi.org/10.1016/j.cognition.2007.05.006

Lewis, G., Solomyak, O., & Marantz, A. (2011). The neural basis of obligatory decomposition of suffixed words. Brain and Language, 118(3), 118–127. doi:https://doi.org/10.1016/j.bandl.2011.04.004

Matchin, W., Brodbeck, C., Hammerly, C., & Lau, E. (2019). The temporal dynamics of structure and content in sentence comprehension: Evidence from fMRI-constrained MEG. Human Brain Mapping, 40(2), 663–678. doi:10.1002/hbm.24403

Matchin, W., Liao, C.-H., Gaston, P., & Lau, E. (2019). Same words, different structures: An fMRI investigation of argument relations and the angular gyrus. Neuropsychologia, 125, 116–128. doi:https://doi.org/10.1016/j.neuropsychologia.2019.01.019

Mollica, F., Siegelman, M., Diachek, E., Piantadosi, S. T., Mineroff, Z., Futrell, R., … Fedorenko, E. (2020). Composition is the Core Driver of the Language-selective Network. Neurobiology of Language, 1(1), 104–134. doi:10.1162/nol_a_00005

Nobre, A. C., Coull, J. T., Frith, C. D., & Mesulam, M. M. (1999). Orbitofrontal cortex is activated during breaches of expectation in tasks of visual attention. Nature Neuroscience, 2(1), 11–12. doi:10.1038/4513

Nolte, G. (2003). The magnetic lead field theorem in the quasi-static approximation and its use for magnetoencephalography forward calculation in realistic volume conductors. Physics in Medicine & Biology, 48(22), 3637.

Payne, B. R., Lee, C. L., & Federmeier, K. D. (2015). Revisiting the incremental effects of context on word processing: Evidence from single-word event-related brain potentials. Psychophysiology, 52(11), 1456–1469. doi:10.1111/psyp.12515

Peelen, M. V., & Caramazza, A. (2012). Conceptual Object Representations in Human Anterior Temporal Cortex. The Journal of Neuroscience, 32(45), 15728–15736. doi:10.1523/jneurosci.1953-12.2012

Penolazzi, B., Hauk, O., & Pulvermuller, F. (2007). Early semantic context integration and lexical access as revealed by event-related brain potentials. Biological Psychology, 74(3), 374–388. doi:10.1016/j.biopsycho.2006.09.008

Pickering, M. J., & Gambi, C. (2018). Predicting While Comprehending Language: A Theory and Review. Psychological Bulletin, 144(10), 1002–1044. doi:10.1037/bul0000158

Price, C. J., & Devlin, J. T. (2011). The Interactive Account of ventral occipitotemporal contributions to reading. Trends in Cognitive Sciences, 15(6), 246–253. doi:https://doi.org/10.1016/j.tics.2011.04.001

Pulvermüller, F., Shtyrov, Y., & Hauk, O. (2009). Understanding in an instant: Neurophysiological evidence for mechanistic language circuits in the brain. Brain and Language, 110(2), 81–94. doi:https://doi.org/10.1016/j.bandl.2008.12.001

Pylkkänen, L. (2008). Mismatching Meanings in Brain and Behavior. Language and Linguistics Compass, 2(4), 712–738. doi:https://doi.org/10.1111/j.1749-818X.2008.00073.x

Pylkkänen, L. (2019). The neural basis of combinatory syntax and semantics. Science, 366(6461), 62–66. doi:10.1126/science.aax0050

Pylkkänen, L. (2020). Neural basis of basic composition: what we have learned from the red&#x2013;boat studies and their extensions. Philosophical Transactions of the Royal Society B: Biological Sciences, 375(1791), 20190299. doi:doi:10.1098/rstb.2019.0299

Pylkkänen, L., Martin, A. E., McElree, B., & Smart, A. (2009). The Anterior Midline Field: Coercion or decision making? Brain and Language, 108(3), 184–190. doi:https://doi.org/10.1016/j.bandl.2008.06.006

Pylkkänen, L., & McElree, B. (2007). An MEG Study of Silent Meaning. Journal of Cognitive Neuroscience, 19(11), 1905–1921. doi:10.1162/jocn.2007.19.11.1905

Ralph, M. A. L., Jefferies, E., Patterson, K., & Rogers, T. T. (2017). The neural and computational bases of semantic cognition. Nature Reviews Neuroscience, 18, 42. doi:10.1038/nrn.2016.150 https://www.nature.com/articles/nrn.2016.150#supplementary-information

Ramnani, N., & Owen, A. M. (2004). Anterior prefrontal cortex: insights into function from anatomy and neuroimaging. Nature Reviews Neuroscience, 5(3), 184–194. doi:10.1038/nrn1343

Rayner, K. (1986). Eye movements and the perceptual span in beginning and skilled readers. Journal of Experimental Child Psychology, 41(2), 211–236. doi:10.1016/0022-0965(86)90037-8

Rayner, K., & Duffy, S. A. (1986). Lexical complexity and fixation times in reading: Effects of word frequency, verb complexity, and lexical ambiguity. Memory & Cognition, 14(3), 191–201.

Rayner, K., & Well, A. D. (1996). Effects of contextual constraint on eye movements in reading: A further examination. Psychonomic Bulletin & Review, 3(4), 504–509.

Reichle, E. D., Pollatsek, A., & Rayner, K. (2012). Using E-Z Reader to simulate eye movements in nonreading tasks: A unified framework for understanding the eye–mind link. Psychological Review, 119(1), 155–185. doi:10.1037/a0026473

Reichle, E. D., Rayner, K., & Pollatsek, A. (2003). The E-Z Reader model of eye-movement control in reading: Comparisons to other models. Behavioral and Brain Sciences, 26(4), 445–476. doi:10.1017/S0140525X03000104

Rice, G. E., Lambon Ralph, M. A., & Hoffman, P. (2015). The Roles of Left Versus Right Anterior Temporal Lobes in Conceptual Knowledge: An ALE Meta-analysis of 97 Functional Neuroimaging Studies. Cerebral Cortex, 25(11), 4374–4391. doi:10.1093/cercor/bhv024

Rubenstein, H., Garfield, L., & Millikan, J. A. (1970). Homographic entries in the internal lexicon. Journal of Verbal Learning and Verbal Behavior, 9(5), 487–494. doi:https://doi.org/10.1016/S0022-5371(70)80091-3

Schäfer, R., & Bildhauer, F. (2012). Building large corpora from the web using a new efficient tool chain. Paper presented at the LREC.

Schoffelen, J.-M., Hultén, A., Lam, N., Marquand, A. F., Uddén, J., & Hagoort, P. (2017). Frequency-specific directed interactions in the human brain network for language. Proceedings of the National Academy of Sciences, 114(30), 8083–8088. doi:10.1073/pnas.1703155114

Schoffelen, J.-M., Oostenveld, R., Lam, N. H. L., Uddén, J., Hultén, A., & Hagoort, P. (2019). A 204-subject multimodal neuroimaging dataset to study language processing. Scientific Data, 6(1), 17. doi:10.1038/s41597-019-0020-y

Schuster, S., Hawelka, S., Himmelstoss, N. A., Richlan, F., & Hutzler, F. (2020). The neural correlates of word position and lexical predictability during sentence reading: evidence from fixation-related fMRI. Language, Cognition and Neuroscience, 35(5), 613–624. doi:10.1080/23273798.2019.1575970

Sereno, S. C., Brewer, C. C., & O’Donnell, P. J. (2003). Context Effects in Word Recognition:Evidence for Early Interactive Processing. Psychological Science, 14(4), 328–333. doi:10.1111/1467-9280.14471

Sereno, S. C., Hand, C. J., Shahid, A., Mackenzie, I. G., & Leuthold, H. (2019). Early EEG correlates of word frequency and contextual predictability in reading. Language Cognition and Neuroscience. doi:10.1080/23273798.2019.1580753

Sereno, S. C., & Rayner, K. (2003). Measuring word recognition in reading: eye movements and event-related potentials. Trends in Cognitive Sciences, 7(11), 489–493. doi:https://doi.org/10.1016/j.tics.2003.09.010

Sharoh, D., van Mourik, T., Bains, L. J., Segaert, K., Weber, K., Hagoort, P., & Norris, D. G. (2019). Laminar specific fMRI reveals directed interactions in distributed networks during language processing. Proceedings of the National Academy of Sciences, 116(42), 21185–21190. doi:10.1073/pnas.1907858116

Smith, M. E., & Halgren, E. (1987). Event-related potentials during lexical decision: effects of repetition, word frequency, pronounceability, and concreteness. Electroencephalogr Clin Neurophysiol Suppl, 40, 417–421.

Staub, A. (2015). The Effect of Lexical Predictability on Eye Movements in Reading: Critical Review and Theoretical Interpretation. Language and Linguistics Compass, 9(8), 311–327. doi:10.1111/lnc3.12151

Staub, A., & Benatar, A. (2013). Individual differences in fixation duration distributions in reading. Psychonomic Bulletin & Review, 20(6), 1304–1311. doi:10.3758/s13423-013-0444-x

Staub, A., & Goddard, K. (2019). The Role of Preview Validity in Predictability and Frequency Effects on Eye Movements in Reading. Journal of Experimental Psychology-Learning Memory and Cognition, 45(1), 110–127. doi:10.1037/xlm0000561

Staub, A., Grant, M., Astheimer, L., & Cohen, A. (2015). The influence of cloze probability and item constraint on cloze task response time. Journal of Memory and Language, 82, 1–17. doi:10.1016/j.jml.2015.02.004

Strijkers, K., Bertrand, D., & Grainger, J. (2015). Seeing the same words differently: the time course of automaticity and top-down intention in reading. J Cogn Neurosci, 27(8), 1542–1551. doi:10.1162/jocn_a_00797

Thacker, J. M., Chambers, C. G., & Graham, S. A. (2018). When it is apt to adapt: Flexible reasoning guides children’s use of talker identity and disfluency cues. Journal of Experimental Child Psychology, 167, 314–327. doi:https://doi.org/10.1016/j.jecp.2017.11.008

Van Den Bosch, A., & Berck, P. (2009). Memory-Based Machine Translation and Language Modeling. Prague Bull. Math. Linguistics, 91, 17–26.

Van Petten, C., & Kutas, M. (1990). Interactions between sentence context and word frequencyinevent-related brainpotentials. Memory & Cognition, 18(4), 380–393. doi:10.3758/bf03197127

Van Veen, B. D., van Drongelen, W., Yuchtman, M., & Suzuki, A. (1997). Localization of brain electrical activity via linearly constrained minimum variance spatial filtering. IEEE Transactions on Biomedical Engineering, 44(9), 867–880. doi:10.1109/10.623056

Veldre, A., Yu, L., Andrews, S., & Reichle, E. D. (2020). Towards a complete model of reading: Simulating lexical decision, word naming, and sentence reading with Über-Reader. Proceedings of the 42nd Annual Conference of the Cognitive Science Society, 151–157. Retrieved from https://hdl.handle.net/2123/22990

Willems, R. M., Frank, S. L., Nijhof, A. D., Hagoort, P., & van den Bosch, A. (2016). Prediction During Natural Language Comprehension. Cerebral Cortex, 26(6), 2506–2516. doi:10.1093/cercor/bhv075

Woollams, A. M. (2015). Lexical is as lexical does: computational approaches to lexical representation. Language, Cognition and Neuroscience, 30(4), 395–408. doi:10.1080/23273798.2015.1005637

